# Diversity of ACE2 and its interaction with SARS-CoV-2 receptor binding domain

**DOI:** 10.1101/2020.10.25.354548

**Authors:** Jessie Low-Gan, Ruiqi Huang, Gabrielle Warner, Abigail Kelley, Duncan McGregor, Vaughn Smider

## Abstract

COVID-19, the clinical syndrome caused by the SARS-CoV-2 virus, has rapidly spread globally causing tens of millions of infections and over a million deaths. The potential animal reservoirs for SARS-CoV-2 are currently unknown, however sequence analysis has provided plausible potential candidate species. SARS-CoV-2 binds to the angiotensin I converting enzyme 2 (ACE2) to enable its entry into host cells and establish infection. We analyzed the binding surface of ACE2 from several important animal species to begin to understand the parameters for the ACE2 recognition by the SARS-CoV-2 spike protein receptor binding domain (RBD). We employed Shannon entropy analysis to determine the variability of ACE2 across its sequence and particularly in its RBD interacting region, and assessed differences between various species’ ACE2 and human ACE2. As cattle are a known reservoir for coronaviruses with previous human zoonotic transfer, and has a relatively divergent ACE2 sequence, we compared the binding kinetics of bovine and human ACE2 to SARS-CoV-2 RBD. This revealed a nanomolar binding affinity for bovine ACE2 but an approximate ten-fold reduction of binding compared to human ACE2. Since cows have been experimentally infected by SARS-CoV-2, this lower affinity sets a threshold for sequences with lower homology to human ACE2 to be able to serve as a productive viral receptor for SARS-CoV-2.

## Introduction

COVID-19, caused by the novel coronavirus SARS-CoV-2, is a zoonotic disease(1–3) that has thus far resulted in over one million deaths worldwide and over 42 million infections (4). The virus crossed the species boundary, possibly from bats and potentially through an intermediate species, to humans and has spread through the respiratory route across the globe in the past year. SARS-CoV-2 is a member of the betacoronavirus genera, which includes other coronaviruses like SARS-CoV, that caused severe acute respiratory syndrome in a pandemic in 2002-2004 (resulting in approximately 800 deaths in 37 countries), and MERS-CoV which caused Middle Eastern Respiratory Syndrome in 2012 (resulting in over 800 deaths in 27 countries). Both of these viruses originated in other species, SARS-CoV in bats, and MERS-CoV in camels, and crossed the species barrier to humans. Other coronaviruses like HCoV-NL63, HCoV-229E, HCoV-HKU1, and HCoV-O43 are known to cause mild respiratory disease in humans and also crossed the species barrier(5–7). Interestingly, the latter two viruses which cause more clinically mild disease appear to have derived from bovine coronavirus (BCoV)(8–10), which causes respiratory and intestinal disease in cattle (7,11–13). Betacoronaviruses as a group have a wide host range including several agricultural and companion animal species(8). Thus, coronavirus disease in humans appears to largely be through zoonotic transfer and has resulted in enormous worldwide morbidity and mortality as well as significant economic loss.

Given that coronaviruses appear to have crossed the species barrier several times in human history with devastating consequences, there is a need to understand the interactions between coronaviruses and their receptors in different animal species. This is particularly important since humans interact with companion animals as well as many agricultural species, often in close quarters where respiratory spread can easily occur. In addition to the impact of spread from species to species on human health, coronaviruses can cause devastating effects to animals resulting in morbidity, mortality, and major economic losses (5,7,11,14). Indeed SARS-CoV-2 has been documented to infect dogs (*Canis lupus*), cats (*Felis catus*) (15), and mink (*Mustela lutrola*) (16) in nature, and ferrets (*Mustela putoris)* (15), hamsters (*Cricetulus griseus*) (17) and cows (*Bos taurus*) (18) have been experimentally infected. Human to tiger transfer occurred at the Bronx Zoo (19,20). A mink farm in the Netherlands was ravaged by SARS-CoV-2 infection and farms in the U.S. and Spain were also recently infected (16,21). Additionally, SARS-CoV-2 is now studied *in vivo* in hamsters (*Cricetulus griseus*) and ferrets (*Mustela putoris*) as animal models to understand viral pathology and evaluate therapeutics. The need to understand animal susceptibility to coronavirus infection is therefore important to public health, the economy, as well as to establish well understood animal models for therapeutic and vaccine development.

SARS-CoV-2 utilizes its trimeric spike protein to bind to the angiotensin I converting enzyme 2 (ACE2) on target pneumocytes or other host cells (22–28). This interaction occurs with high affinity, and results in viral membrane fusion to the host cell and initiates the infectious process. SARS-CoV-1 also utilizes ACE2 as a receptor, however their spike proteins bind with lower affinity (31 nM *K*_D_) than SARS-CoV-2 (4.2 nM *K*_D_)(23). Crystal structure and electron microscopy analysis of SARS-CoV-2 with ACE2 has revealed the interacting amino acid residues of the spike receptor binding domain (RBD) and the human ACE2 surface(23,25,29,30). With sequences available for many companion and agriculturally important species, the ability to assess potential spike RBD binding is an important first step towards prediction of infection of these alternative hosts. Here we analyze the conservation and diversity of the ACE2 protein in multiple important animal species, with particular emphasis on the region that interacts with SARS-CoV-2 spike RBD. We confirm that an ACE2 from *Bos taurus*, which is somewhat divergent from human ACE2 in its binding region, interacts with SARS-CoV-2 spike RBD with high affinity, suggesting that multiple mammalian species may be susceptible to infection with this coronavirus.

## Results

In order to assess the potential of SARS-CoV-2 to interact with ACE2 of important companion and agricultural species, we assembled the ACE2 sequences of several species (Table 1 and Supplemental Table 1). The human ACE2/RBD cocrystal structure (PDB: 6M17)(27) was utilized to visualize interacting residues between the SARS-CoV-2 spike RBD and human ACE2 (Figure 1). In ACE2, 8 residues on two helices make direct contact with spike RBD and an additional 17 residues are within 5 angstroms of the RBD. These 25 residues are color coded in Figure 1 as purple (contact) and cyan (nearby), respectively. We focused our evaluation on these interacting residues in the following analyses.

**Table 1.** Various species ACE2 percent identity to human ACE2. The percent identity was calculated across all amino acids of ACE2 (Full length ACE2) or only the interacting residues as shown in Figure 1.

|  |  | % Sequence Identity to Human ACE2 |  |
| --- | --- | --- | --- |
|  | Species | Full length ACE2 | Interacting Residues |
| Human | *Homo sapiens* | 100 | 100 |
| Cynomolgus monkey | *Macaca fascicularis* | 95.03 | 100 |
| Rhesus monkey | *Macaca mulatta* | 94.78 | 100 |
| Horse | *Equus caball* | 86.71 | 72 |
| Donkey | *Equus asinus* | 85.57 | 72 |
| Cat | *Felis catus* | 85.22 | 80 |
| Rabbit | *Oryctolagus cuniculus* | 84.84 | 76 |
| Pangolin | *Manis javanica* | 84.72 | 68 |
| Golden hamster | *Mesocricetus auratus* | 84.47 | 88 |
| Chinese hamster | *Cricetulus griseus* | 84.35 | 88 |
| Dog | *Canis lupus* | 83.33 | 76 |
| Ferret | *Mustela putorius* | 82.48 | 68 |
| Mouse | *Mus musculus* | 82.11 | 68 |
| Goat | *Capra hircus* | 81.72 | 84 |
| Sheep | *Ovis aries* | 81.69 | 84 |
| Pig | *Sus scrofa* | 81.37 | 76 |
| Bat | *Rhinolophus sinicus* | 80.75 | 72 |
| Cow | *Bos taurus* | 78.8 | 84 |
| Chicken | *Gallus gallus* | 66.25 | 52 |

**Figure 1.**
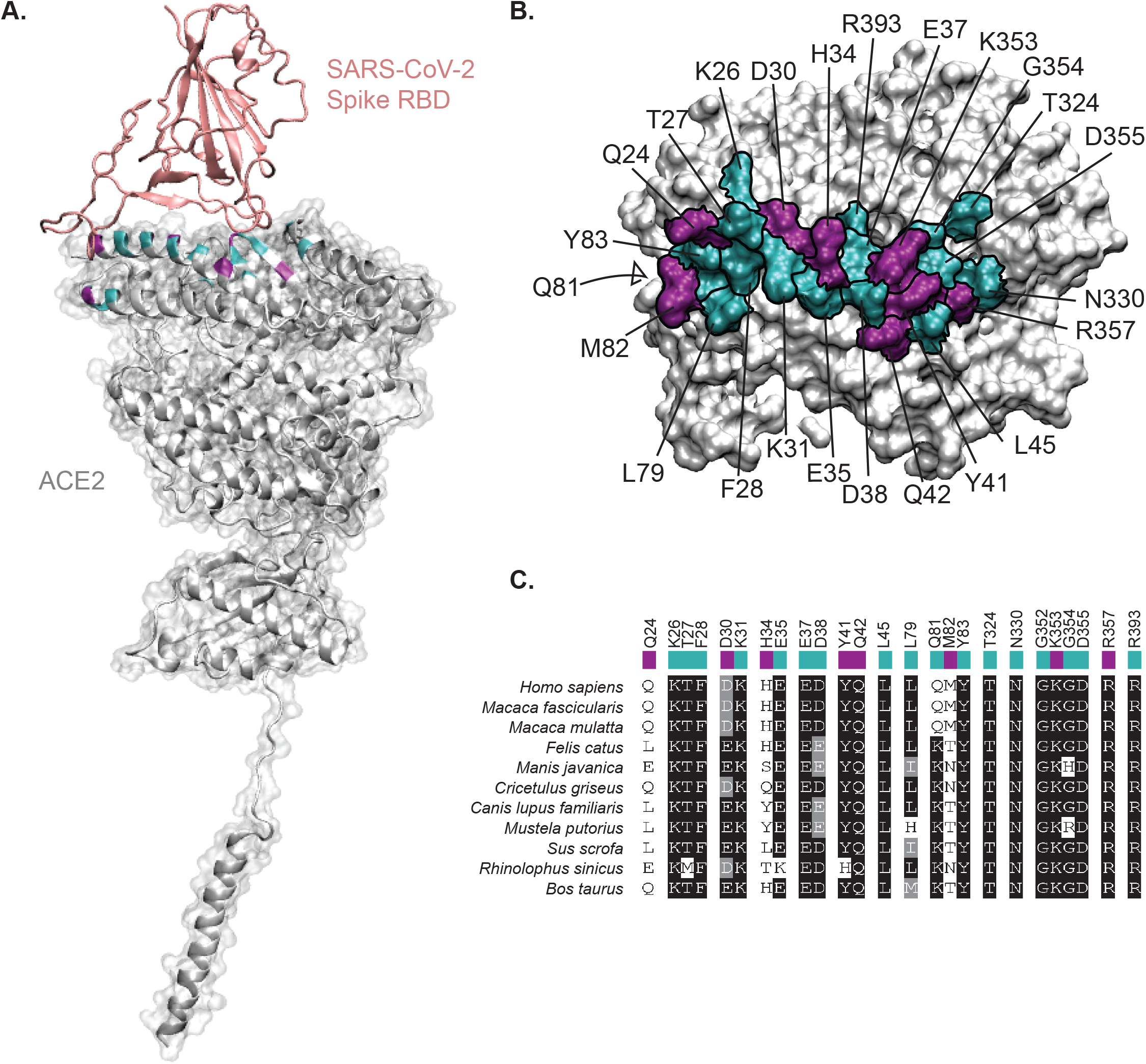
ACE2 residues interacting with SARS-CoV-2 spike RBD. **(A).** The cocrystal structure for the spike RBD (salmon color) and ACE2 (grey, PDB: 6M17) was used to visualize residues that contact the RBD (shaded purple, Q24, D30, H34, Y41, Q42, M82, K353, R357) or are within 5 A of the RBD (shaded cyan, K26, T27, F28, K31, K35, E37, D38, L45, L79, Q81, Y83, T324, N330, G352, G354, , D355, R393). Throughout the manuscript we refer to these as “contact residues” (purple) and “nearby residues” (cyan), and together as “interacting residues”. **(B)** Top down view of ACE2 in space filling mode, with residues color-coded as in (A). G352 is not visible in this orientation. **(C)** Boxshade alignment of only the contact residues and nearby amino acid residues from (B) for multiple species. Contact residues are indicated with a cyan bar on top of the sequence, and nearby residues with a purple bar.

In order to determine the overall differences between ACE2 of different species, we employed protein sequence alignment as well as variability analysis (Supplemental Figures 1 and 2). All vertebrate ACE2 sequences showed significant homology to human ACE2, with cynomologous and rhesus monkeys being 95% and 94% identical to human (Table 1). Other mammals were between 80-87% identical to human. Horseshoe bats, a potential reservoir for SARS-CoV-2 (28,31) was only 81% and 76% identical through the entire sequence and interacting residues, respectively. Surprisingly, the percent identity across the entire ACE2 sequence did not fully correlate with percent identity of the interacting residues. For example, cows show lower homology throughout the entire ACE2 sequence at only 79% compared to the other species, but is 84% identical within the identified interacting residues. In contrast, dogs are 83.3% identical across the entire sequence, but only 76% identical in the interacting residues. A similar lower identity in binding site interacting residues is also seen for rabbits (Table 1). The relative contribution of the interacting residues versus residues outside the RBD binding site is currently not known, although it is expected that the interacting residues are far more important to infection relative to the residues outside of the RBD binding site.

For variability analysis, the structural importance of protein regions, and even individual amino acid residues, can be compared across multiple species. Such diversity analyses initially identified the complementary determining regions within antibodies as important interacting domains with antigen by Kabat and Wu(32,33), and more recently Shannon entropy evaluation has been employed to identify conserved and diverse domains of multiple proteins through multiple sequence analysis(34,35). First, we aligned the ACE2 sequences and determined their percent identities (Table 1 and Supplemental Table 1). Then, we calculated Shannon entropy (SE) across the ACE2 sequences. Of note, ACE2 is remarkably conserved across its sequence, with few residues exhibiting high variability (Figure 2).

**Figure 2.**
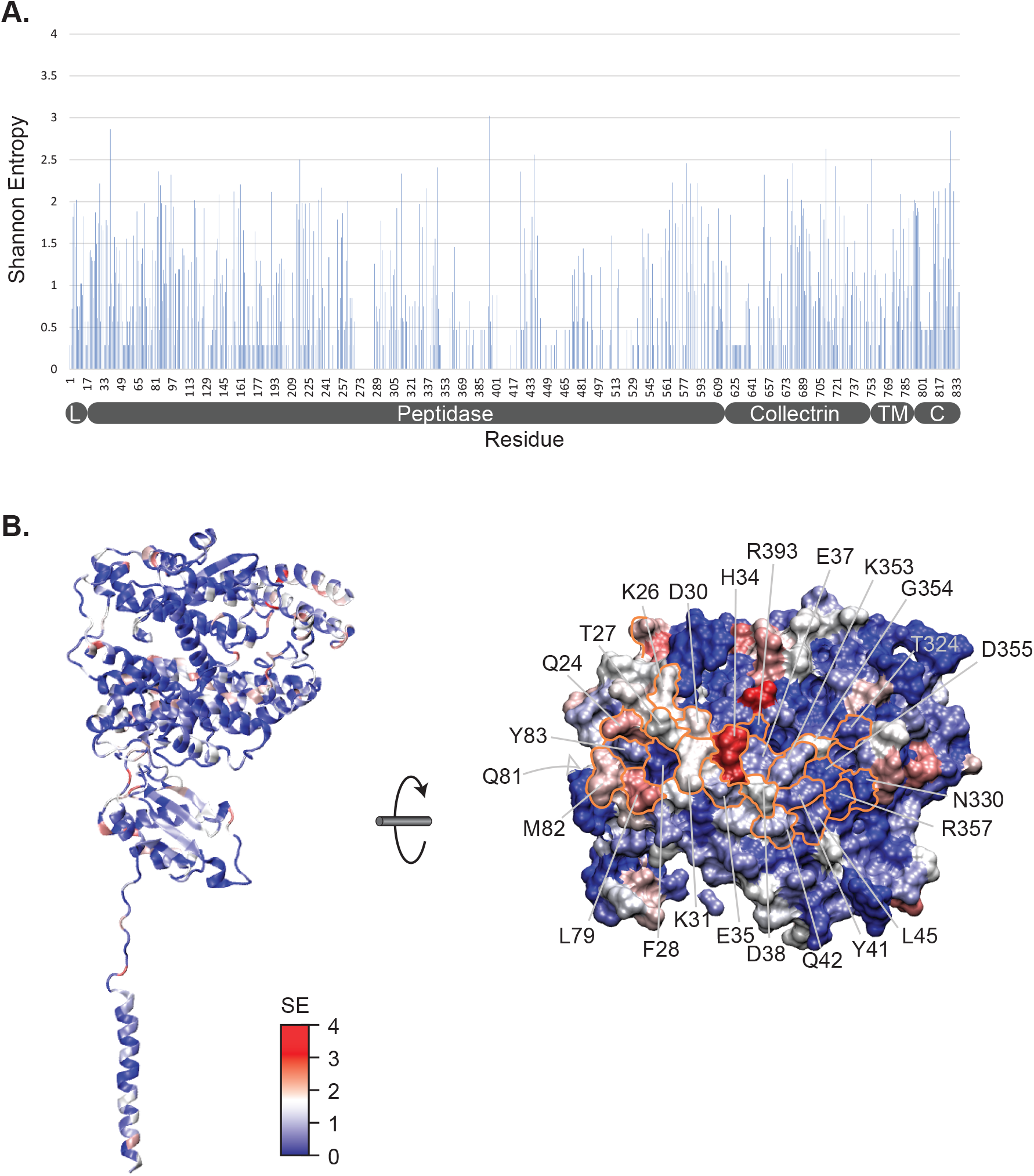
Shannon entropy analysis of ACE2 shows low variability. **(A)** Shannon entropy plot of the entire ACE2 sequence. The residue positions, and schematic of functional domains are illustrated below the plot. Few residues have values above 2.5, indicating low variability **(B)** (Left) Shannon entropy values projected onto the human ACE2 structure as a heat map from blue (low) to red (high). (Right) View of the RBD-interacting surface as in Figure 1, with contact and nearby residues labeled. Blue residues are very highly conserved.

Within the interacting residues, 21 of 25 are highly conserved, with SE values below 2. Significantly, no residues had values above 3. The more variable residues are colored red and conserved residues blue in Figure 2. Five residues, N330, G352, D355, R357, and R393, are completely conserved, with SE values of zero (Figure 2 and Supplemental Table 1). Four of these are nearby residues with RBD, with only R357 being a contact residue. The complete conservation of these residues suggests that they play an important role in the protease function or structural integrity of ACE2. Residues with values between 0 and 1 are K353, which is a contact residue, and L45, Y83, T324, which are nearby residues (Supplemental Table 1). Amino acids with SE values between 1-2 are K26, T27, D30, K31, E35, D38, Q42, M82, and G354. The most diverse residues, with SE values over 2, are Q24, H34, L79, and Q81. Of these, Q24 and H34 are contact residues, with H34 found in a central location in the ACE2-RBD interface (Figure 1, B), and having by far the highest SE value at 2.88. Others have analyzed the evolution of ACE2 residues and have found positions 24 and 34 to be undergoing positive evolutionary selection pressure (36), and suggested that these positions could play a role in predicting infectivity by SARS-CoV-2 (36).

Since the interacting residues are likely most important for viral interaction with ACE2 on the host cell, we evaluated the residues that differed between the various species’ ACE2 and human ACE2 in this region (Figure 3 and Supplemental Figure 3). As mentioned, horseshoe bat (*Rhinolophus sinicus*) shares only 19/25 interacting residues (76%) with human ACE2, and only 5/8 contact residues. Specifically, D30E, H34T, Y41H, and M82N (contact residues), and T27M, E35K (nearby residues) are mutated in horseshoe bat ACE2 relative to human ACE2 (Figure 3, upper left), suggesting potentially lower affinity for spike RBD. Pangolin, a possible intermediate host of SARS-CoV-2, has only one contact residue (M82N) and three nearby residues (D38E, L79I, and G354H) altered, and both D38E and L79I are conservative changes (Figure 3, upper right). Felines, which have had documented natural infection (15,20), have only two active site residues mutated (D38E and M82T), and only M82T is a contact residue (Figure 3, middle left). Dogs, which also have been infected naturally, have similar binding site residues as cat but with the notable exception of H34Y (Figure 3, middle), a residue reported to be important in binding. Cow has three interaction site residues mutated, of which M82T is a contact residue, and pig has 5 mutations where Q24L, D30E, H34L, M82T are contact residues (Figure 3 bottom).

**Figure 3.**
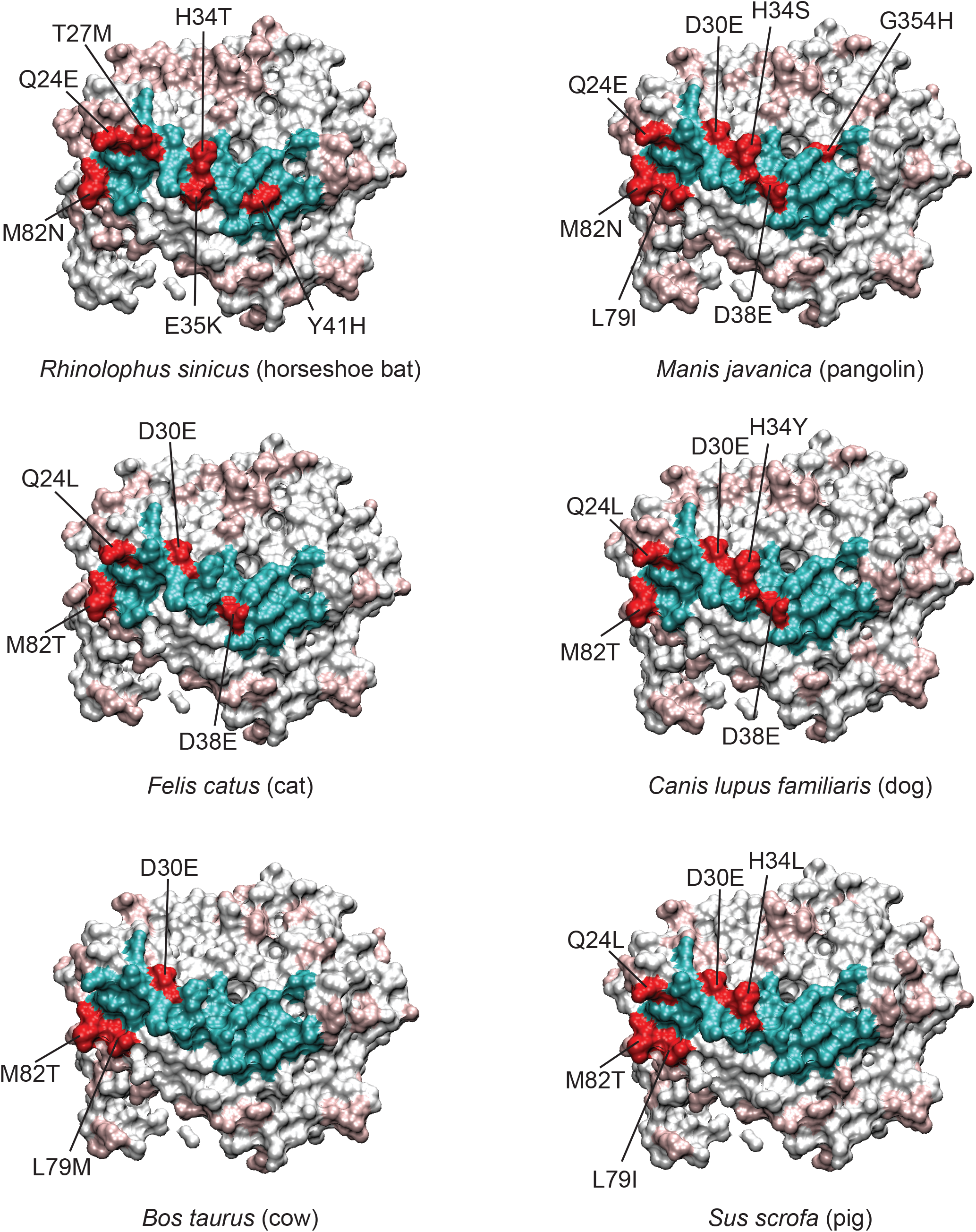
Variable residues at the ACE2-RBD interface of individual species. With human ACE2 as a reference, the variant interacting residues for each species are colored red, and conserved residues colored cyan. For the remainder of the ACE2 protein, conserved residues are white and variable residues light red. Certain residues like M82, Q24, D30, and H34 are often mutated relative to human. D30 is conservatively changed to glutamate, however more non-conservative changes can be seen for H34, for example. More species have been analyzed and are shown in Supplemental Figure 3.

Cows serve as a reservoir for bovine coronavirus (BCoV) a respiratory infection of cattle that is a betacoronavirus (11,12) distantly related to SARS-CoV-2(7). Of considerable note, two BCoV-related coronaviruses, HCoV-OC43 and HCoV-HKU1, have crossed the species barrier, with OC43 likely from cows to humans to cause “common cold” respiratory disease in humans(37). Whereas BCoV, HCoV-OC43 and HCoV-HKU1 utilize 9-*O*-acetylated sialoglycans as cellular receptors(38), and SARS-CoV-2 utilizes ACE2, cows are a potential important species to evaluate for possible SARS-CoV-2 infection as they are a known coronavirus reservoir and they could theoretically serve as host for SARS-CoV-2 and other coronaviruses like BCoV, which could potentially provide a host for coronavirus recombination, selection, and evolution. From a biochemical standpoint, cows have a somewhat more distantly related ACE2 protein compared to many other vertebrates (78.8% compared to most other species which are over 80%), however their binding site residues are more conserved (84%)(Table 1). Therefore, it would be useful to know whether the lower homology across the entire ACE2 sequence prohibits productive ACE2/RBD interaction. To address this question, we expressed human and bovine ACE2 as antibody Fc fusion proteins and compared their interaction with SARS-CoV-2 by enzyme linked immunosorbent assay (ELISA) and further quantified their binding kinetics by surface plasmon resonance analysis (Figure 4). By ELISA, bovine ACE2 had an approximately ten-fold worse binding EC_50_ (0.129 nM for human ACE2 compared to 1.299 nM for bovine ACE2)(Figure 4, A). This lower apparent affinity for bovine ACE2 was confirmed by surface plasmon resonance analysis which showed a *K*_D_ for bovine ACE2 of 36.25 nM versus 7.5 nM for human ACE2 (Figure 4, B and Supplementary Table 1). This dissociation constant difference relates primarily to a faster off-rate for bovine ACE2 compared to human ACE2 (Supplementary Figure 5). Of note, despite this lower *K*_D_ for bovine ACE2, this affinity is very similar to the *K*_D_ for human ACE2 for the RBD of SARS-CoV-1. Interestingly, for both human and bovine ACE2, the SPR data fit more consistently with a two-site model for interaction, suggesting that the RBD may be multimerizing to produce avidity effects on the chip surface. A potential second site would have ten-fold lower *K*_D_ values (Supplemental Figure 5), which interestingly, are more in line with the EC_50_ values of the ELISA (Figure 4, A). Such interactions would be important to explore to understand the details of interaction between coronavirus spike RBD and ACE2 on the cell surface. These interactions would be important to inhibit by therapeutic agents, for example by monoclonal antibodies targeting the virus. The avidity interaction may decrease the *K*_D_ (increase the affinity) of bovine ACE2 with spike RBD from 36 nM to 2.5 nM, and 7.5 nM to 0.4 nM for human ACE2, a substantial enhancement of the interaction. Regardless of the mechanisms of interaction, it is clear that bovine ACE2 still has high affinity towards SARS-CoV-2 RBD, albeit with five to ten fold worse binding than human ACE2, but yet can still mediate infection of bovine cells(18).

**Figure 4.**
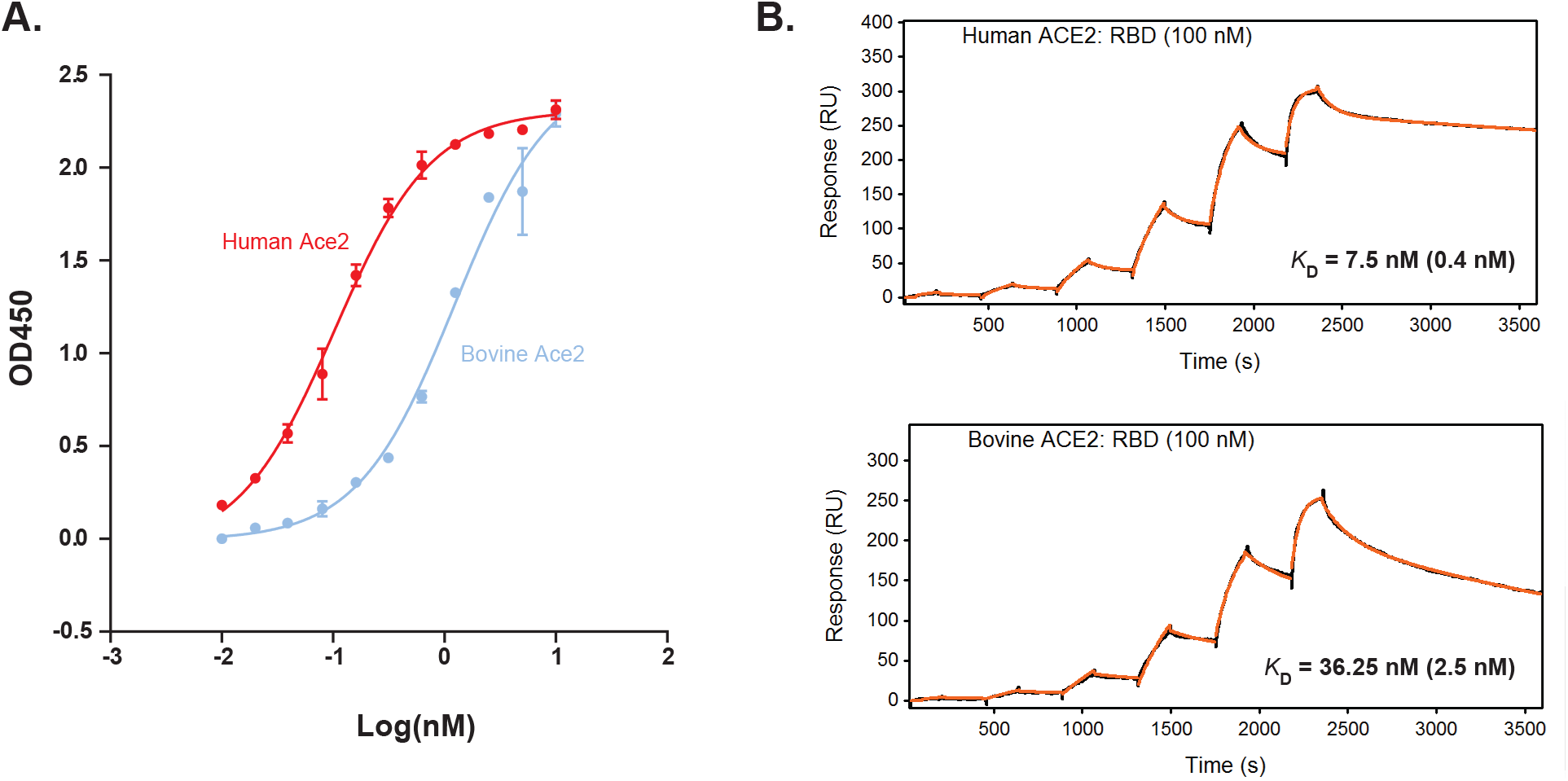
SARS-CoV-2 spike RBD binds to bovine ACE2 with lower affinity than human ACE2. **(A)** ELISA of human (red) and bovine (blue) ACE2-Fc binding to immobilized SARS-CoV-2 spike RBD. **(B)** Surface plasmon resonance analysis of RBD as an analyte on immobilized Human ACE-2 Fc (top) or bovine ACE2-Fc (bottom). The data was fit to a two-site model (Supplemental Figure 5). The *K*_D_ for site one is indicated, and site two is in parentheses.

## Discussion

The recent COVID-19 pandemic has spread rapidly across the globe through human populations, but additionally has also infected several animal species. While zoonotic in origin, it is still unclear which species provided the reservoir for transfer to humans. Coronaviruses as a group have a very wide range of host species, and have jumped the species barrier multiple times to humans(39,40). While bats appear to be a host species for coronaviruses related to SARS-CoV and SARS-CoV-2, it is possible that an as yet unidentified species serves as the reservoir for this virus. Additionally, it is clear that SARS-CoV-2 can naturally infect other species, such as cats, dogs, and mink. In order to understand the host range of SARS-CoV-2 as well as identify potential reservoirs for the virus it is critically important to understand (i) details about the identity and binding properties between the virus and its host cell receptor and possible co-receptors, (ii) other biological requirements needed for the virus to replicate and transmit, for example host cell enzymes needed for viral processing, replication, and assembly. For SARS-CoV-2, ACE2 appears to be the major receptor required for cell entry, so understanding its interaction with ACE2 from humans as well as other species is important to enable predictive methods for viral host range. Here we analyze ACE2 diversity across several species, and specifically evaluate binding of SARS-CoV-2 RBD to bovine ACE2, finding key interacting residues to be highly conserved.

Several studies have used sequence homology and/or structural modeling to attempt to predict SARS-CoV-2 RBD binding to various species’ ACE2 in order to predict the virus host range(15,36,39–43). In an effort to predict species permissive to infection, Damas et.al. developed a five-tiered scoring scheme based on percent identity of 410 vertebrate species, as well as specific structural features of SARS-CoV-1 or SARS-CoV-2 interactions with ACE2(36). They also focused on 25 amino acid residues at the RBD binding interface, however their 25 residues differed from ours in that they included S16, N53, N90, and N322, with the asparagines included as potential glycosylation sites that may impact RBD binding. However, our approach was agnostic in choosing residues that were either (i) known contact residues with the RBD, or (ii) within 5 angstroms of contact residues. Residues included in our analysis which were not included in Damas et.al. were K26, Q81, and N352. Damas et. al. also note that the host range of SARS-CoV-2 might be quite broad and suggest new species that should be evaluated for animal models of virus infection, which notably include cows, which scored in their “medium” category for predictive binding to RBD. Like our assessment of homology, they find that bats score very low in predicted ACE2 binding. Interestingly, using sequence evolution and selection analysis, they identify Q24 and H34 as positions undergoing positive selection and evolution. We identified these as amongst the most variable residues by Shannon entropy analysis, and these also appear to be important residues at the ACE2/RBD interface. As in our analysis, pangolins scored low in potential RBD binding based on interacting residue homology, suggesting that SARS-CoV-2 may bind other pangolin receptors, or have other mechanisms to interact with ACE2. As pangolins are thought to be a possible intermediate host for SARS-CoV-2, much more biochemical and infectivity data with this controversial species should be obtained.

In a different approach, Lam et.al. used structural modeling to predict the change in free energy, ΔΔG, for 215 ACE2 sequences derived from different species(44). They correlated the ΔΔG with published infectability information to provide a framework to predict which species may be susceptible to SARS-CoV-2 infection. Like other studies, their work suggests a broad range of mammal susceptibility, with the exception of non-placental mammals. They similarly find that horseshoe bats have higher ΔΔG values (*i.e.* lower affinity), calling into question their susceptibility and potential as a reservoir for SARS-CoV-2.

There have been several studies that have measured the *K*_D_ between SARS-CoV-2 spike (or RBD) and ACE2, with values ranging from 1.2 to 44 nM (23,25,29,30). In these studies, differences in the experimental conditions, such as whether ACE2 or RBD was immobilized, and technique used such as biolayer interferometry versus surface plasmon resonance, could account for differences in kinetic values. In our study, we immobilized ACE2 and measured RBD interaction as the analyte using surface plasmon resonance, which is similar to Lan et. al. who reported a value of *K*_D_ = 4.6 nM, which is close to our value of 7.5 nM. However, Lan et.al. as well as all of the other studies utilized a 1:1 model for binding, whereas we found evidence for a two site complexation on the surface, and applied a two site model that gave *K*_D1_ and *K*_D2_ values of 7.5 nM and 0.4 nM, respectively. Of considerable note, Forssen et.al. reanalyzed the binding data from Lan et.al. and Tian et.al. and found evidence of multimeric interactions, similar to our study(45). Since the RBD exists in close proximity to two other RBD subunits in the spike protein trimer, it is plausible that cooperative interactions between RBDs exist during ACE2 interaction events. Such events could considerably enhance the interactions providing avidity to enable infection. In this regard, ACE2 variants from species with mutations in the RBD interface may still be permissive to infection because of the cooperative and enhanced avidity of binding events.

One of the major challenges in developing predictive methods for viral infectivity is the limited amount of both infection and RBD-ACE2 biochemical interaction data for most species. For example, only few species have had documented infection in the real world, or even in well-controlled experimental systems(15,36,40,44). In this regard, detailed biochemical analysis of multiple species’ receptors would provide valuable information that could be used in predictive modeling studies. With the decreased costs of synthetic DNA and high-throughput screening approaches, such analyses could potentially be accomplished rapidly. As a first step in this process, here we evaluate cow ACE2 for binding SARS-CoV-2 RBD by both ELISA and surface plasmon resonance, a species with documented experimental infection by SARS-CoV-2 and of significant agricultural importance that serves as a reservoir for other betacoronaviruses, including at least one, HCoV-O43, that has jumped the species barrier to become an endemic respiratory virus in humans. The affinity of bovine ACE2 is nearly ten-fold worse than human ACE2 in binding the coronavirus RBD, however binding is still in the mid-nanomolar range. Of potential importance is the possible cooperativity in binding that may occur between ACE2 and RBD, which we observed in surface plasmon resonance analysis which suggested a two-site model for binding. Interestingly, this data was consistent with ELISA binding data, suggesting *K*_D_ in the subnanomolar range for human, and low nanomolar range for bovine ACE2. Further study into various species’ viral receptors, including the recently discovered neuropilin-2 co-receptor for SARS-CoV-2(46), should enable much more accurate and rapid development of predictive algorithms based on viral receptor sequence and structural modelling.

## Methods

### Sequence and structure analysis

ACE2 sequences from 19 species were obtained from the NCBI protein sequence database. A number of multiple sequence alignments were performed using Clustal Omega. Specific multiple sequence alignments for (i) companion animals (*Canis lupus familiaris, Felis catus, Mus musculus, Mesocricetus auratus, Oryctolagus cuniculus*), (ii) agriculturally important animals (*Bos taurus, Capra hircus, Equus asinus, Equus caballus, Gallus gallus, Ovis aries, and Sus scrofa*), and (iii) species relevant to animal models or potential reservoirs of interest to vaccine testing research or suspected reservoirs (*Macaca mulatta, Macaca fascicularis, Felis catus, Manis javanica, Cricetulus griseus, Canis lupus familiaris, Mustela putoris, Sus scrofa, Rhinolophus sinicus, Bos taurus*) were investigated. In all sequence alignments the *Homo sapiens* ACE2 sequence was included as a reference.

Further analysis on the multiple sequence alignments were carried out with a variety of software. Phylogenetic trees and pairwise percent identities were obtained with Clustal Omega (47) and visualized in UGENE (http://ugene.net). The open-source EMBL Boxshade Server was used to produce the boxshade alignments (https://embnet.vital-it.ch/software/BOX_form.html). The Protein Variability Server (http://imed.med.ucm.es/PVS/) was employed to calculate Shannon Entropy values for each residue position.

For structural analysis, the ACE2 and SARS-CoV-2 Receptor Binding Domain (RBD) complex structure files (PDB) were obtained from the RCSB PDB (PDB ID: 6M17)(27). Eight contact residues of ACE2 were identified based on the co-crystal structure (27), and we additionally added 17 ACE2 residues (nearby residues) that were within at least 5 angstroms of the RBD region and classified the full list as “interaction residues”. Thus, there were a total of 25 interaction residues, with 8 known RBD contact residues and 17 nearby residues. Using a *de novo* python script, these residues were extracted into a separate “sequence” and an additional, specific multiple sequence alignment was constructed.

The interaction residue multiple sequence alignments were investigated to give more focused insight into which species would be at risk of infection, with the assumption that residues directly interacting with the SARS-CoV-2 spike protein would be essential for infection. Structural analyses were performed in Visual Molecular Dynamics (VMD, https://www.ks.uiuc.edu/Research/vmd/).

### Protein purification

Human and bovine ACE2 proteins were produced as fusion proteins to human IgG1 Fc according to our published methods for monoclonal antibody purification[2-4]. Briefly, 30 M HEK293 Freestyle cells were transfected with 293fectin combined with 30 μg of pFuse-based vectors containing the human or bovine ACE2 fused to the human Fc region. Cells were shaken at 37 °C for 4 days with 8% CO_2_. The media was clarified by centrifugation at 4000 RPM for 5 minutes followed by filtration through a 0.22 μm filter. The media was concentrated and buffer-exchanged into PBS using Amicon Ultra Centrifugal Filter unit (MWCO = 10,000) (MilliporeSigma) at 4 °C. The concentrated media was then loaded onto a protein A-sepharose column (Cytiva) pre-equilibrated with 20 mM sodium phosphate, pH 7.0, followed by washing of the column with 10 column volumes of the same buffer and eluted twice with 1 column volume of 0.1 M glycine-HCl, pH 2.7 into fractions containing 0.1 column volume of 1M Tris, pH 8. Purified proteins were buffer-exchanged into PBS using Amicon Ultra Centrifugal Filter unit (MWCO = 10,000) (MilliporeSigma), quantified using 280 nM absorbance on a Nanodrop spectrophotometer (Thermo Fisher Scientific), and resolved on an SDS-PAGE stained with InstantBlue Coomassie Protein Stain (Abcam).

The SARS-CoV-2 RBD in plasmid NR-52309 (BEI Resources) was transfected and harvested as described above, but purified using TALON cobalt metal affinity resin (Takara Bio) following the manufacturer’s protocol, except that 50 mM, 100 mM, 200 mM and 300 mM imidazole gradient elution fractions (1 column volume of each) were collected. Each elution fraction was resolved on an SDS-PAGE gel stained with InstantBlue Coomassie Protein Stain (Abcam), and fractions containing a single RBD band were pooled, buffer-exchanged into PBS and quantified as described above. Purified protein was resolved on SDS-PAGE with Coomassie (Gel-code blue) staining.

### Enzyme linked immunosorbent assay (ELISA)

ELISA plates (Corning 3690) were coated with 100 ng soluble SARS-CoV-2 RBD at 4 °C in 1 × PBS (pH 7.4) overnight. Plates were washed three times with Tris buffered saline pH 7.4 containing 0.1% Tween-20 (TBST) and blocked with 2% milk (Marvel, dried skim milk dissolved in TBST) at room temperature for 1 h. Serially diluted human or bovine ACE2-Fc were added to the wells in 2% milk/TBST, plates were incubated at room temperature for 1 h, washed four times with TBST and then goat anti-human Fc-HRP (Jackson ImmunoResearch #109-035-098, diluted 1: 5000 in 2% milk/TBST) was added to the wells. Plates were incubated at room temperature for 30 minutes and washed five times with TBST, then developed by adding 50 μl TMB substrate solution (Thermo Scientific) per well and incubated at room temperature for 3 minutes. The HRP-TMB reaction was stopped by adding 50 μl 1.0 N sulfuric acid per well. The optical density at 450 nm was read on a microplate reader (SPECTRAMAX M2, Molecular Devices). Antigen-binding curves and EC_50_ values were generated and calculated using four parameter logistic regression in GraphPad Prism 8 (GraphPad Software, Inc., San Diego, CA).

### Surface plasmon resonance

Surface plasmon resonance of immobilized ACE2 proteins to RBD as the analyte was performed by Biosensor Tools (https://www.biosensortools.com/, Salt Lake City, Utah). ACE2-Fc ligands were captured onto a Protein A surface at four different surface densities. Data were collected in Hepes Buffered Saline (HBS) containing 0.1 mg/ml BSA at 25 degrees C. The RBD analyte (36 kDa) was tested in a 3-fold titration series up to 1 uM and 100 nM for both ACE2 protein surfaces at the 4 different densities. The data for each data set were fit to a two independent site model. The *K*_D_ values indicated in Figure 4 are the average of four experiments at different Rmax values for 100 nM analyte.

---

**Supplemental Table 1. Shannon Entropy for RBD interacting residues of ACE2.**

**Supplemental Figure 1. Boxshade amino acid sequence alignment of multiple ACE2 species.**

**Supplemental Figure 2. Phylogenetic tree of ACE2 from multiple species.**

**Supplemental Figure 3. Structures showing variations of the ACE2-RBD interface for four species. This analysis was identical to Figure 3.**

**Supplemental Figure 4. Pairwise ACE2 percent identities for multiple vertebrate species.**

**Supplemental Figure 5. Surface plasmon resonance data for human and bovine ACE2-Fc binding SARS-CoV-2 RBD.**

## Acknowledgements

The following reagent was produced under HHSN272201400008C and obtained through BEI Resources, NIAID, NIH: Vector pCAGGS Containing the SARS-Related Coronavirus 2, Wuhan-Hu-1 Spike Glycoprotein Receptor Binding Domain (RBD), NR-52309. We thank David Myszka of Biosensor Tools for surface plasmon resonance data analysis.

## Author Contributions

JG and VS designed experiments and analyzed data. JG assembled sequences and performed bioinformatic analysis and generated figures. RH designed SARS-CoV-2 RBD expression protocols and purified the RBD protein. GW and AK purified ACE2-Fc proteins and performed ELISA assays and analyzed SPR results. DM and GW designed ACE2 expression vectors. VS and JG wrote the manuscript. VS supervised the project.

