## Supplemental Data for "Diversity of ACE2 and its interaction with SARS-CoV-2 receptor binding domain"

**Supplemental Table 1. Shannon Entropy values for RBD interacting residues.** Contact residues are in purple and nearby residues are in cyan.

| Residue | Shannon Entropy |
|---|---|
| Q24 | 2.215 |
| K26 | 1.717 |
| T27 | 1.657 |
| F28 | 0.286 |
| D30 | 1.782 |
| K31 | 1.717 |
| H34 | 2.866 |
| E35 | 1.022 |
| E37 | 0.748 |
| D38 | 1.578 |
| Y41 | 0.569 |
| Q42 | 1.022 |
| L45 | 0.569 |
| L79 | 2.361 |
| Q81 | 2.195 |
| M82 | 1.98 |
| Y83 | 0.811 |
| T324 | 0.469 |
| N330 | 0 |
| G352 | 0 |
| K353 | 0.922 |
| G354 | 1.457 |
| D355 | 0 |
| R357 | 0 |
| R393 | 0 |

```
Homo_sapiens     1 MSSSSWLLLSLVAVTAAQSTIEEQAKTFLDKFNHEAEDLFYQSSLASWNYNTNITEENVQ
Felis_catus_     1 MSGSFWLLLSFAALTAAQSTTEELAKTFLEKFNHEAEELSYQSSLASWNYNTNITDENVQ
Oryctolagus_     1 MSGSSWLLLSLVAVTAAQSTIEELAKTFLEKFNQEAEDLSYQSALASWDYNTNITEENVQ
Mesocricetus     1 MSSSSWLLLSLVAVTTAQSIIEEQAKTFLDKFNQEAEDLSYQSALASWNYNTNITEENAQ
Canis_lupus_     1 MSGSSWLLLSLAALTAAQS-TEDLVKTFLEKFNYEAEELSYQSSLASWNYNINITDENVQ
Mus_musculus     1 MSSSSWLLLSLVAVTTAQSLTEENAKTFLNNFNQEAEDLSYQSSLASWNYNTNITEENAQ
Equus_caball     1 MSGSSWLLLSLVAVTAAQSTTEDLAKTFLEKFNSEAEELSHQSSLASWSYNTNITDENVQ
Equus_asinus     1 MSGSSWLLLSLVAVTAAQSTTEDLAKTFLEKFNSEAEELSHQSSLASWSYNTNITDENVQ
Capra_hircus     1 MTGSFWLLLSLVAVTAAQSTTEEQAKTFLEKFNHEAEDLSYQSSLASWNYNTNITDENVQ
Ovis_aries__     1 MTGSFWLLLSLVAVTAAQSTTEGQAKTFLEKFNHEAEDLSYQSSLASWNYNTNITDENVQ
Sus_scrofa__     1 MSGSFWLLLSLIPVTAAQSTTEELAKTFLEKFNLEAEDLAYQSSLASWTINTNITDENIQ
Bos_taurus__     1 MTGSFWLLLSLVAVTAAQSTTEEQAKTFLEKFNHEAEDLSYQSSLASWNYNTNITDENVQ
Gallus_gallu     1 MLLHFWLLCGLSAVVTPQDVT-QEAQTFLAEFNVRAEDISYENSLASWNYNTNITEETAR
Macaca_fasci     1 MSGSSWLLLSLVAVTAAQSTIEEQAKTFLDKFNHEAEDLFYQSSLASWNYNTNITEENVQ
Macaca_mulat     1 MSGSSWLLLSLVAVTAAQSTIEEQAKTFLDKFNHEAEDLFYQSSLASWNYNTNITEENVQ
Manis_javani     1 MSGSSWLLLSLVAVTAAQSTSDEEAKTFLEKFNSEAEELSYQSSLASWNYNTNITDENVQ
Cricetulus_g     1 MSSSSWLLLSLVAVTTAQSIIEEQAKTFLDKFNQEAEDLSYQSALASWNYNTNITEENAQ
Mustela_puto     1 MLGSSWLLLSLAALTAAQSTTEDLAKTFLEKFNYEAEELSYQNSLASWNYNTNITDENIQ
Rhinolophus_     1 MSGSSWLLLSLVAVTTAQSTTEDEAKMFLDKFNTKAEDLSHQSSLASWDYNTNINDENVQ

Homo_sapiens    61 NMNNAGDKWSAFLKEQSTLAQMYPLQEIQNLTVKLQLQALQQNGSSVLSEDKSKRLNTIL
Felis_catus_    61 KMNEAGAKWSAFYEEQSKLAKTYPLAEIHNTTVKRQLQALQQSGSSVLSADKSQRLNTIL
Oryctolagus_    61 KMNDAEAKWSAFYEEQSKLAKTYPSQEVQNLTVKRQLQALQQSGSSALSADKSKQLNTIL
Mesocricetus    61 KMNEAAAKWSAFYEEQSKLAKNYSLQEVQNLTIKRQLQALQQSGSSALSADKNKQLNTIL
Canis_lupus_    60 KMNNAGAKWSAFYEEQSKLAKTYPLEEIQDSTVKRQLRALQHSGSSVLSADKNQRLNTIL
Mus_musculus    61 KMSEAAAKWSAFYEEQSKTAQSFSLQEIQTPIIKRQLQALQQSGSSALSADKNKQLNTIL
Equus_caball    61 KMNEAGARWSAFYEEQCKLAKTYPLEEIQNLTVKRQLQALQQSGSSVLSADKSKRLNEIL
Equus_asinus    61 KMNEAGARWSAFYEEQCKLAKTYPLEEIQNLTVKRQLQALQQSGSSVLSADKSKRLNEIL
Capra_hircus    61 KMNEARAKWSAFYEEQSRMARTYSLEEIQNLTLKRQLKALQHSGTSVLSAEKSKRLNTIL
Ovis_aries__    61 KMNEARAKWSAFYEEQSRMARTYSLEEIQNLTLKRQLKALQHSGTSVLSAEKSKRLNTIL
Sus_scrofa__    61 KMNDARAKWSAFYEEQSRIAKTYPLDEIQTLILKRQLQALQQSGTSGLSADKSKRLNTIL
Bos_taurus__    61 KMNEARAKWSAFYEEQSRMAKTYSLEEIQNLTLKRQLKALQHSGTSALSAEKSKRLNTIL
Gallus_gallu    60 KMSEAGAKWAAFYEEASRNASRFSLANIQDAVTRLQIQSLQDRGSSVLSPEKYSRLNSVM
Macaca_fasci    61 NMNNAGEKWSAFLKEQSTLAQMYPLQEIQNLTVKLQLQALQQNGSSVLSEDKSKRLNTIL
Macaca_mulat    61 NMNNAGEKWSAFLKEQSTLAQMYPLQEIQNLTVKLQLQALQQNGSSVLSEDKSKRLNTIL
Manis_javani    61 KMNVAGAKWSTFYEEQSKIAKNYQLQNIQNDTIKRQLQALQLSGSSALSADKNQRLNTIL
Cricetulus_g    61 KMNEAAAKWSAFYEEQSKLAKNYSLQEVQNLIIKRQLQALQQSGSSALSADKNKQLNTIL
Mustela_puto    61 KMNIAGAKWSAFYEEESQHAKTYPLEEIQDPIIKRQLRALQQSGSSVLSADKRERLNTIL
Rhinolophus_    61 KMDEAGAKWSAFYEEQSKLAKNYSLEQIQNVTVKLQLQILQQSGSPVLSEDKSKRLNSIL

Homo_sapiens   121 NTMSTIYSTGKVCNPDNPQECLLLEPGLNEIMANSLDYNERLWAWESWRSEVGKQLRPLY
Felis_catus_   121 NAMSTIYSTGKACNPNNPQECLLLEPGLDDIMENSKDYNERLWAWEGWRAEVGKQLRPLY
Oryctolagus_   121 STMSTIYSTGKVCNQSNPQECFLLEPGLDEIMAKSTDYNERLWAWEGWRSVVGKQLRPLY
Mesocricetus   121 NTMSTIYSTGKVCNPKNPQECLLLEPGLDDIMATSTDYNERLWAWEGWRAEVGKQLRPLY
Canis_lupus_   120 NSMSTVYSTGKACNPSNPQECLLLEPGLDDIMENSKDYNERLWAWEGWRSEVGKQLRPLY
Mus_musculus   121 NTMSTIYSTGKVCNPKNPQECLLLEPGLDEIMATSTDYNSRLWAWEGWRAEVGKQLRPLY
Equus_caball   121 NTMSTIYSTGKVCNPSNPQECLLLEPGLDAIMENSKDYNQRLWAWEGWRSEVGKQLRPLY
Equus_asinus   121 NTMSTIYSTGKVCNPSNPQECLLLEPGLDAIMENSKDYNQRLWAWEGWRSEVGKQLRPLY
Capra_hircus   121 NKMSTIYSTGKVLDPN-TQECLALEPGLDDIMENSRDYNRRLWAWEGWRAEVGKQLRPLY
Ovis_aries__   121 NKMSTIYSTGKVLDPN-TQECLALEPGLDDIMENSRDYNRRLWAWEGWRAEVGKQLRPLY
Sus_scrofa__   121 NTMSTIYSSGKVLDPNNPQECLVLEPGLDEIMENSKDYSRRLWAWESWRAEVGKQLRPLY
Bos_taurus__   121 NKMSTIYSTGKVLDPN-TQECLALEPGLDDIMENSRDYNRRLWAWEGWRAEVGKQLRPLY
Gallus_gallu   120 NSMSTIYSTGVVCKATEPFDCLVLEPGLDDIMANSIDYHERLWAWEGWRADVGRMMRPLY
Macaca_fasci   121 NTMSTIYSTGKVCNPNNPQECLLLDPGLNEIMEKSLDYNERLWAWEGWRSEVGKQLRPLY
Macaca_mulat   121 NTMSTIYSTGKVCNPNNPQECLLLDPGLNEIMEKSLDYNERLWAWEGWRSEVGKQLRPLY
Manis_javani   121 NTMSTIYSTGKVCNPGNPQECSLLEPGLDNIMESSKDYNERLWAWEGWRSEVGKQLRPLY
Cricetulus_g   121 NTMSTIYSTGKVCNPKNPQECLLLEPGLDDIMATSTDYNERLWAWEGWRAEVGKQLRPLY
Mustela_puto   121 NAMSTIYSTGKACNPNNPQECLLLEPGLDDIMENSKDYNERLWAWEGWRSEVGKQLRPLY
Rhinolophus_   121 NAMSTIYSTGKVCKPNKPQECLLLEPGLDNIMGTSKDYNERLWAWEGWRAEVGKQLRPLY
```

```
Homo_sapiens  181 EEYVVLKNEMARANHYEDYGDYWRGDYEVNGVDGYDYSRGQLIEDVEHTFEEIKPLYEHL
Felis_catus_  181 EEYVALKNEMARANNYEDYGDYWRGDYEEEWTDGYNYSRSQLIKDVEHTFTQIKPLYQHL
Oryctolagus_  181 EEYVVLKNEMARANNYEDYGDYWRADYEAEGADGYDYSRSQLIDDVERTFSEIKPLYEQL
Mesocricetus  181 EEYVVLKNEMARANNYEDYGDYWRGDYEAEGADGYNYNGNQLIEDVERTFKEIKPLYEQL
Canis_lupus_  180 EEYVALKNEMARANNYEDYGDYWRGDYEEEWENGYNYSRNQLIDDVELTFTQIMPLYQHL
Mus_musculus  181 EEYVVLKNEMARANNYNDYGDYWRGDYEAEGADGYNYNRNQLIEDVERTFAEIKPLYEHL
Equus_caball  181 EEYVVLKNEMARANNYEDYGDYWRGDYEAEGPSGYDYSRDQLIEDVERTFAEIKPLYEHL
Equus_asinus  181 EEYVVLKNEMARANNYEDYGDYWRGDYEAEGPSGYDYSRDQLIEDVERTFAEVTEEV---
Capra_hircus  180 EEYVVLENEMARANNYEDYGDYWRGDYEVTGAGDYDYSRDQLMKDVERTFAEIKPLYEQL
Ovis_aries__  180 EEYVVLENEMARANNYEDYGDYWRGDYEVTGAGDYDYSRDQLMKDVERTFEEIKPLYEQL
Sus_scrofa__  181 EEYVVLENEMARANNYEDYGDYWRGDYEVTGTGDYDYSRNQLMEDVERTFAEIKPLYEHL
Bos_taurus__  180 EEYVVLENEMARANNYEDYGDYWRGDYEVTGAGDYDYSRDQLMKDVERTFAEIKPLYEQL
Gallus_gallu  180 EEYVELKNEAARLNNYSDYGDYWRANYETDYPEEYKYSRDQLVQDVEKTFEQIKPLYQHL
Macaca_fasci  181 EEYVVLKNEMARANHYKDYGDYWRGNYEVNGVDGYDYNRDQLIEDVERTFEEIKPLYEHL
Macaca_mulat  181 EEYVVLKNEMAGANHYKDYGDYWRGDYEVNGVDGYDNNRDQLIEDVERTFEEIKPLYEHL
Manis_javani  181 EEYVVLKNEMARANHYEDYGDYWRGDYEAEGANGYNYSRDHLIEDVEHIFTQIKPLYEHL
Cricetulus_g  181 EEYVVLKNEMARANNYKDYGDYWRGDYEAEGADGYNYNGNQLIEDVERTFKEIKPLYEQL
Mustela_puto  181 EEYVALKNEMARANNYEDYGDYWRGDYEEEWADGYSYSRNQLIEDVEHTFTQIKPLYEHL
Rhinolophus_  181 EEYVVLKNEMARGYHYEDYGDYWRRDYETEESPGPGYSRDQLMKDVERIFTEIKPLYEHL

Homo_sapiens  241 HAYVRAKLMNAYPS-YISPIGCLPAHLLGDMWGRFWTNLYSLTVPFGQKPNIDVTDAMVD
Felis_catus_  241 HAYVRAKLMDTYPS-RISPTGCLPAHLLGDMWGRFWTNLYPLTVPFGQKPNIDVTDAMVN
Oryctolagus_  241 HAYVRTKLMDAYPS-RISPTGCLPAHLLGDMWGRFWTNLYSLTVPFGQKPNIDVTDTMVN
Mesocricetus  241 HAYVRTKLMNTYPS-YISPTGCLPAHLLGDMWGRFWTNLYPLTVPFGQKPNIDVTDAMVN
Canis_lupus_  240 HAYVRTKLMDTYPS-YISPTGCLPAHLLGDMWGRFWTNLYPLTVPFGQKPNIDVTNAMVN
Mus_musculus  241 HAYVRRKLMDTYPS-YISPTGCLPAHLLGDMWGRFWTNLYPLTVPFAQKPNIDVTDAMMN
Equus_caball  241 HAYVRAKLMDTYPS-HINPTGCLPAHLLGDMWGRFWTNLYSLTVPFGQKPNIDVTDAMVD
Equus_asinus  238 -----------YPD-T--------DVGQCDMWGRFWTNLYSLTVPFGQKPNIDVTDAMVD
Capra_hircus  240 HAYVRAKLMNTYPS-YISPTGCLPAHLLGDMWGRFWTNLYSLTVPFEHKPSIDVTEKMKN
Ovis_aries__  240 HAYVRAKLMDTYPS-YISPTGCLPAHLLGDMWGRFWTNLYSLTVPFEHKPSIDVTEKMKN
Sus_scrofa__  241 HAYVRAKLMDAYPS-RISPTGCLPAHLLGDMWGRFWTNLYPLTVPFGEKPSIDVTEAMVN
Bos_taurus__  240 HAYVRAKLMHTYPS-YISPTGCLPAHLLGDMWGRFWTNLYSLTVPFEHKPSIDVTEKMEN
Gallus_gallu  240 HAYVRHRLEQVYGSELINPTGCLPAHLLGDMWGRFWTNLYNLTVPYPEKPNIDVTSAMAQ
Macaca_fasci  241 HAYVRAKLMNAYPS-YISPTGCLPAHLLGDMWGRFWTNLYSLTVPFGQKPNIDVTDAMVN
Macaca_mulat  241 HAYVRAKLMNAYPS-YISPTGCLPAHLLGDMWGRFWTNLYSLTVPFGQKPNIDVTDAMVN
Manis_javani  241 HAYVRAKLMDNYPS-HISPTGCLPAHLLGDMWGRFWTNLYPLTVPFRQKPNIDVTDAMVN
Cricetulus_g  241 HAYVRTKLMDTYPS-YISPTGCLPAHLLGDMWGRFWTNLYPLTVPFGQKPNIDVTDAMVN
Mustela_puto  241 HAYVRAKLMDAYPS-RISPTGCLPAHLLGDMWGRFWTNLYPLMVPFRQKPNIDVTDAMVN
Rhinolophus_  241 HAYVRAKLMDTYPF-HISPTGCLPAHLLGDMWGRFWTNLYPLTVPFGQKPNIDVTDEMLK

Homo_sapiens  300 QAWDAQRIFKEAEKFFVSVGLPNMTQGFWENSMLTDPGNVQKAVCHPTAWDLGKGDFRIL
Felis_catus_  300 QSWDARRIFKEAEKFFVSVGLPNMTQGFWENSMLTEPGDSRKVVCHPTAWDLGKGDFRIK
Oryctolagus_  300 QGWDAERIFKEAEKFFVSVGLPSMTHGFWENSMLPESGDGRKVVCHPTAWDLGKRDFRIK
Mesocricetus  300 QGWNAERIFKEAEKFFVSVGLPYMTQGFWENSMLTDPGDDRKVVCHPTAWDLGKGDFRIK
Canis_lupus_  299 QSWDARKIFKEAEKFFVSVGLPNMTQEFWGNSMLTEPSDSRKVVCHPTAWDLGKGDFRIK
Mus_musculus  300 QGWDAERIFQEAEKFFVSVGLPHMTQGFWANSMLTEPADGRKVVCHPTAWDLGHGDFRIK
Equus_caball  300 QSWDAKRIFEEAEKFFVSVGLPNMTQGFWENSMLTEPGDGRKVVCHPTAWDLGKGDFRIK
Equus_asinus  278 QSWDAKRIFEEAEKFFVSVGLPNMTQGFWENSMLTEPGDGRKVVCHPTAWDLGKGDFRIK
Capra_hircus  299 QSWDAERIFKEAEKFFVSIGLPYMTQGFWNNSMLTEPGDGRKVVCHPTAWDLGKGDFRIK
Ovis_aries__  299 QSWDAERIFKEAEKFFVSIGLPYMTQGFWDNSMLTEPGDGRKVVCHPTAWDLGKGDFRIK
Sus_scrofa__  300 QSWDAIRIFEEAEKFFVSIGLPNMTQGFWNNSMLTEPGDGRKVVCHPTAWDLGKGDFRIK
Bos_taurus__  299 QSWDAERIFKEAEKFFVSISLPYMTQGFWDNSMLTEPGDGRKVVCHPTAWDLGKGDFRIK
Gallus_gallu  300 KNWDAMKIFKTAEAFFASIGLYNMTEGFWTNSMLTEPTDNRKVVCHPTAWDMGKNDYRIK
Macaca_fasci  300 QAWNAQRIFKEAEKFFVSVGLPNMTQGFWENSMLTDPGNVQKVVCHPTAWDLGKGDFRII
Macaca_mulat  300 QAWNAQRIFKEAEKFFVSVGLPNMTQGFWENSMLTDPGNVQKVVCHPTAWDLGKGDFRII
Manis_javani  300 QTWDANRIFKEAEKFFVSVGLPKMTQTFWENSMLTEPGDGRKVVCHPTAWDLGKHDFRIK
Cricetulus_g  300 QGWDAERIFKEAEKFFVSVGLPHMTQGFWGNSMLTDPGDDRKVVCHPTAWDLGKGDFRIK
Mustela_puto  300 QSWDARRIFEEAETFFVSVGLPNMTEGFWQNSMLTEPGDNRKVVCHPTAWDLGKRDFRIK
Rhinolophus_  300 QGWDADRIFKEAEKFFVSVGLPNMTEGFWNNSMLTEPGDGRKVVCHPTAWDLGKGDFRIK
```

```
Homo_sapiens  360 MCTKVTMDDFLTAHHEMGHIQYDMAYAAQPFLLRNGANEGFHEAVGEIMSLSAATPKHLK
Felis_catus_  360 MCTKVTMDDFLTAHHEMGHIQYDMAYAVQPFLLRNGANEGFHEAVGEIMSLSAATPNHLK
Oryctolagus_  360 MCTKVTMDNFLTAHHEMGHIQYDMAYATQPFLLRNGANEGFHEAVGEIMSLSAATPEHLK
Mesocricetus  360 MCTKVTMDNFLTAHHEMGHIQYDMAYATQPFLLRNGANEGFHEAVGEIMSLSAATPEHLK
Canis_lupus_  359 MCTKVTMDDFLTAHHEMGHIQYDMAYAAQPFLLRNGANEGFHEAVGEIMSLSAATPNHLK
Mus_musculus  360 MCTKVTMDNFLTAHHEMGHIQYDMAYARQPFLLRNGANEGFHEAVGEIMSLSAATPKHLK
Equus_caball  360 MCTKVTMDDFLTAHHEMGHIQYDMAYAVQPYLLRNGANEGFHEAVGEIMSLSAATPNHLK
Equus_asinus  338 MCTKVTMDDFLTAHHEMGHIQYDMAYAVQPYLLRNGANEGFHEAVGEIMSLSAATPNHLK
Capra_hircus  359 MCTKVTMDDFLTAHHEMGHIQYDMAYATQPYLLRNGANEGFHEAVGEIMSLSAATPHYLK
Ovis_aries__  359 MCTKVTMDDFLTAHHEMGHIQYDMAYATQPYLLRNGANEGFHEAVGEIMSLSAATPHYLK
Sus_scrofa__  360 MCTKVTMDDFLTAHHEMGHIQYDMAYAIQPYLLRNGANEGFHEAVGEIMSLSAATPHYLK
Bos_taurus__  359 MCTKVTMDDFLTAHHEMGHIQYDMAYAAQPYLLRNGANEGFHEAVGEIMSLSAATPHYLK
Gallus_gallu  360 MCTKVTMDDFLTAHHEMGHIEYDMAYSVQPFLLRDGANEGFHEAVGEIMSLSAATPQHLK
Macaca_fasci  360 MCTKVTMDDFLTAHHEMGHIQYDMAYAAQPFLLRNGANEGFHEAVGEIMSLSAATPKHLK
Macaca_mulat  360 MCTKVTMDDFLTAHHEMGHIQYDMAYAAQPFLLRNGANEGFHEAVGEIMSLSAATPKHLK
Manis_javani  360 MCTKVTMDDFLTAHHEMGHIQYDMAYAMQPYLLRNGANEGFHEAVGEIMSLSAATPKHLK
Cricetulus_g  360 MCTKVTMDNFLTAHHEMGHIQYDMAYATQPFLLRNGANEGFHEAVGEIMSLSAATPKHLK
Mustela_puto  360 MCTKVTMDDFLTAHHEMGHIQYDMAYAEQPFLLRNGANEGFHEAVGEIMSLSAATPNHLK
Rhinolophus_  360 MCTKVTMEDFLTAHHEMGHIQYDMAYASQPYLLRNGANEGFHEAVGEVMSLSVATPKHLK

Homo_sapiens  420 SIGLLSPDFQEDNETEINFLLKQALTIVGTLPFTYMLEKWRWMVFKGEIPKDQWMKKWWE
Felis_catus_  420 TIGLLSPGFSEDSETEINFLLKQALTIVGTLPFTYMLEKWRWMVFKGEIPKEQWMQKWWE
Oryctolagus_  420 SIGLLPYDFHEDNETEINFLLKQALTIVGTLPFTYMLEKWRWMVFKGEIPKEQWMQKWWE
Mesocricetus  420 SIGLLPSDFQEDNETEINFLLKQALTIVGTLPFTYMLEKWRWMVFKGDIPKEQWMEKWWE
Canis_lupus_  419 NIGLLPPSFFEDSETEINFLLKQALTIVGTLPFTYMLEKWRWMVFKGEIPKDQWMKTWWE
Mus_musculus  420 SIGLLPSDFQEDSETEINFLLKQALTIVGTLPFTYMLEKWRWMVFRGEIPKEQWMKKWWE
Equus_caball  420 AIGLLPPDFYEDSETEINFLLKQALTIVGTLPFTYMLEKWRWMVFKGEIPKEEWMKKWWE
Equus_asinus  398 AIGLLPPDFYEDSETEINFLLKQALTIVGTLPFTYMLEKWRWMVFKGEIPKEEWMKKWWE
Capra_hircus  419 ALGLLAPDFYEDNETEINFLLKQALTIVGTLPFTYMLEKWRWMVFKGEIPKQQWMEKWWE
Ovis_aries__  419 ALGLLAPDFYEDNETEINFLLKQALTIVGTLPFTYMLEKWRWMVFKGEIPKQQWMEKWWE
Sus_scrofa__  420 ALGLLPPDFYEDSETEINFLLKQALTIVGTLPFTYMLEKWRWMVFKGEIPKEQWMQKWWE
Bos_taurus__  419 ALGLLAPDFHEDNETEINFLLKQALTIVGTLPFTYMLEKWRWMVFKGEIPKQQWMEKWWE
Gallus_gallu  420 SLDLLEPTFQEDEETEINFLLKQALTIVGTMPFTYMLEKWRWMVFNGEITKQEWTKRWWK
Macaca_fasci  420 SIGLLSPDFQEDNETEINFLLKQALTIVGTLPFTYMLEKWRWMVFKGEIPKDQWMKKWWE
Macaca_mulat  420 SIGLLSPDFQEDNETEINFLLKQALTIVGTLPFTYMLEKWRWMVFKGEIPKDQWMKKWWE
Manis_javani  420 NIGLLPPDFYEDNETEINFLLKQALTIVGTLPFTYMLEKWRWMVFSGQIPKEQWMKKWWE
Cricetulus_g  420 SIGLLPSNFHEDNETEINFLLKQALTIVGTLPFTYMLEKWRWMVFKGDIPKEKWMEKWWE
Mustela_puto  420 NIGLLPPDFSEDSETDINFLLKQALTIVGTLPFTYMLEKWRWMVFKGEIPKEQWMQKWWE
Rhinolophus_  420 TMGLLSPDFREDNETEINFLLKQALNIVGTLPFTYMLEKWRWMVFKGEIPKEEWMKKWWE

Homo_sapiens  480 MKREIVGVVEPVPHDETYCDPASLFHVSNDYSFIRYYTRTLYQFQFQEALCQAAKHEGPL
Felis_catus_  480 MKREIVGVVEPVPHDETYCDPASLFHVANDYSFIRYYTRTIYQFQFQEALCRIAKHEGPL
Oryctolagus_  480 MKREIVGVVEPMPHDETYCDPAALFHVANDYSFIRYYTRTIYQFQFQEALCQAAQHEGPL
Mesocricetus  480 MKREIVGVVEPLPHDETYCDPAALFHVSNDYSFIRYYTRTIYQFQFQEALCQAAKHDGPL
Canis_lupus_  479 MKRNIVGVVEPVPHDETYCDPASLFHVANDYSFIRYYTRTIYQFQFQEALCQIAKHEGPL
Mus_musculus  480 MKREIVGVVEPLPHDETYCDPASLFHVSNDYSFIRYYTRTIYQFQFQEALCQAAKYNGSL
Equus_caball  480 MKREIVGVVEPVPHDETYCDPAALFHVANDYSFIRYYTRTIYQFQFQEALCQTAKHEGPL
Equus_asinus  458 MKREIVGVVEPVPHDETYCDPAALFHVANDYSFIRYYTRTIYQFQFQEALCQTAKHEGPL
Capra_hircus  479 MKREIVGVVEPLPHDETYCDPACLFHVAEDYSFIRYYTRTIYQFQFHEALCKTAKHEGAL
Ovis_aries__  479 MKREIVGVVEPLPHDETYCDPACLFHVAEDYSFIRYYTRTIYQFQFHEALCKTAKHEGAL
Sus_scrofa__  480 MKREIVGVVEPLPHDETYCDPACLFHVAEDYSFIRYYTRTIYQFQFHEALCRTAKHEGPL
Bos_taurus__  479 MKREIVGVVEPLPHDETYCDPACLFHVAEDYSFIRYYTRTIYQFQFHEALCKTAKHEGAL
Gallus_gallu  480 MKREIVGVVEPVPHDETYCDPAALFHVANDYSFIRYYTRTIYQFQFQEALCKAANHTGPL
Macaca_fasci  480 MKREIVGVVEPVPHDETYCDPASLFHVSNDYSFIRYYTRTLYQFQFQEALCQAAKHEGPL
Macaca_mulat  480 MKREIVGVVEPVPHDETYCDPASLFHVSNDYSFIRYYTRTLYQFQFQEALCQAAKHEGPL
Manis_javani  480 MKREIVGVVEPVPHDETYCDPASLFHVANDYSFIRYYTRTIYQFQFQEALCQTAKHEGPL
Cricetulus_g  480 MKREIVGVVEPLPHDETYCDPAALFHVSNDYSFIRYYTRTIYQFQFQEALCQAAKHDGPL
Mustela_puto  480 MKRDIVGVVEPLPHDETYCDPAALFHVANDYSFIRYYTRTIYQFQFQEALCQIAKHEGPL
Rhinolophus_  480 MKRKIVGVVEPVPHDETYCDPASLFHVANDYSFIRYYTRTIFEFQFHEALCRIAQHDGPL
```

```
Homo_sapiens  540 HKCDISNSTEAGQKLFNMLRLGKSEPWTLALENVVGAKNMNVRPLLNYFEPLFTWLKDQN
Felis_catus_  540 HKCDISNSSEAGKKLLQMLTLGKSKPWTLALEHVVGEKKMNVTPLLKYFEPLFTWLKEQN
Oryctolagus_  540 HKCDISNSTEAGQKLLNMLRLGRSEPWTLALENVVGAKNMDVRPLLNYFEPLFTWLKEQN
Mesocricetus  540 HKCDISNSTEAGQKLLNMLRLGKSEPWTLALENVVGARNMDVRPLLNYFEPLSVWLKEQN
Canis_lupus_  539 HKCDISNSSEAGQKLLEMLKLGKSKPWTYALEIVVGAKNMDVRPLLNYFEPLFTWLKEQN
Mus_musculus  540 HKCDISNSTEAGQKLLKMLSLGNSEPWTKALENVVGARNMDVKPLLNYFQPLFDWLKEQN
Equus_caball  540 HKCDISNSTEAGQKLLQMLSLGKSEPWTLALERIVGVKNMDVRPLLNYFEPLFTWLKDQN
Equus_asinus  518 HKCDISNSTEAGQKLLQMLSLGKSEPWTLALERIVGVKNMDVRPLLNYFEPLFTWLKDQN
Capra_hircus  539 FKCDISNSTEAGQRLLQMLRLGKSEPWTLALENIVGIKTMDVKPLLNYFEPLFTWLKEQN
Ovis_aries__  539 FKCDISNSTEAGQRLLQMLRLGKSEPWTLALENIVGIKTMDVKPLLNYFEPLFTWLKEQN
Sus_scrofa__  540 YKCDISNSTEAGQKLLQMLSLGKSEPWTLALENIVGVKTMDVKPLLSYFEPLLTWLKAQN
Bos_taurus__  539 FKCDISNSTEAGQRLLQMLRLGKSEPWTLALENIVGIKTMDVKPLLNYFEPLFTWLKEQN
Gallus_gallu  540 HKCDITNSTAAGGNLRQLLELGKSKPWTQALESATGEKYMNATPLLHYFEPLFNWLQKNN
Macaca_fasci  540 HKCDISNSTEAGQKLLNMLKLGKSEPWTLALENVVGAKNMNVRPLLNYFEPLFTWLKDQN
Macaca_mulat  540 HKCDISNSTEAGQKLLNMLKLGESEPWTLALENVVGAKNMNVRPLLNYFEPLFTWLKDQN
Manis_javani  540 HKCDISNSAEAGQKLLQMLSLGKSKPWTLALERVVGTKNMDVRPLLNYFEPLLTWLKEQN
Cricetulus_g  540 HKCDISNSTEAGQKLLNMLRLGKSEPWTLALENVVGARNMDVRPLLNYFEPLSVWLKEQN
Mustela_puto  540 YKCDISNSSEAGQKLHEMLSLGRSKPWTFALERVVGAKTMDVRPLLNYFEPLFTWLKEQN
Rhinolophus_  540 HKCDISNSTDAGKKLHQMLSVGKSQAWTKTLEDIVDSRNMDVGPLLKYFEPLYTWLQEQN

Homo_sapiens  600 KNSFVGWSTDWSPYADQSIKVRISLKSALGDRAYEWNDNEMYLFRSSVAYAMRQYFLKVK
Felis_catus_  600 RNSFVGWNTDWRPYADQSIKVRISLKSALGDEAYEWNDNEMYLFRSSVAYAMREYFSKVK
Oryctolagus_  600 RNSFVGWSTEWTPYADQSIKVRISLKTALGDQAYEWNDSEMYLFRSSVAYAMRKYFSEVK
Mesocricetus  600 KNSFVGWNTDWSPYADQSIKVRISLKSALGENAYEWDDNEMYLFRASVAYAMRVYFAKNK
Canis_lupus_  599 RNSFVGWNTDWSPYADQSIKVRISLKSALGEKAYEWNNNEMYLFRSSIAYAMRQYFSEVK
Mus_musculus  600 RNSFVGWNTEWSPYADQSIKVRISLKSALGANAYEWTNNEMFLFRSSVAYAMRKYFSIIK
Equus_caball  600 KNSFVGWSTNWSPYADQSIKVRISLKSALGEKSYEWNDNEMYLFQSSVAYAMRVYFLKAK
Equus_asinus  578 KNSFVGWSTNWSPYADQSIKVRISLKSALGEKAYEWNDNEMYLFQSSVAYAMRVYFLKAK
Capra_hircus  599 RNSFVGWSTEWTPYSDQSIKVRISLKSALGENAYEWNDNEMYLFRSSVAYAMRKYFLEDR
Ovis_aries__  599 RNSFVGWSTEWTPYSDQSIKVRISLKSALGENAYEWNDNEMYLFRSSVAYAMRKYFLKER
Sus_scrofa__  600 GNSSVGWNTDWTPYADQSIKVRISLKSALGEDAYEWNDNEMYLFRSSIAYAMRNYFSSAK
Bos_taurus__  599 RNSFVGWSTEWTPYSDQSIKVRISLKSALGENAYEWNDNEMYLFQSSVAYAMRKYFSEAR
Gallus_gallu  600 SGRSIGWNTDWTPYSDNAIKVRISLKAALGDDAYVWDASELFLFKSSIAYAMRKYFAKEK
Macaca_fasci  600 KNSFVGWSTDWSPYADQSIKVRISLKSALGDKAYEWNDNEMYLFRSSVAYAMRTYFLEIK
Macaca_mulat  600 KNSFVGWSTDWSPYADQSIKVRISLKSALGDKAYEWNDNEMYLFRSSVAYAMRTYFLEIK
Manis_javani  600 KNSFVGWNTDWSPYAAQSIKVRISLKSALGEKAYEWNDSEMYLFRSSVAYAMREYFSKVK
Cricetulus_g  600 KNSFVGWNTDWSPYADQSIKVRISLKSALGENAYEWNDNEMYLFRATVAYAMRVYFAKNK
Mustela_puto  600 RNSFVGWNTDWSPYADQSIKVRISLKSALGEKAYEWNDNEMYFFQSSIAYAMREYFSKVK
Rhinolophus_  600 RKSYVGWNTDWSPYSDQSIKVRISLKSALGENAYEWNDNEMYLFRSSVAYAMREYFLKEK

Homo_sapiens  660 NQMILFGEEDVRVANLKPRISFNFFVTAPKNVSDIIPRTEVEKAIRMSRSRINDAFRLND
Felis_catus_  660 NQTIPFVEDNVWVSNLKPRISFNFFVTASKNVSDVIPRSEVEEAIRMSRSRINDAFRLDD
Oryctolagus_  660 NQTILFGEEDVRVSDLKPRISFNFFVTAPNNVNDIIPRNEVEEAISMSRSRINDIFRLDD
Mesocricetus  660 TQTVPFGVEDIRVSDLKPRVSFNFFVTSPQNVSDIIPRNEVEEAVRLSRGRINDVFGLDD
Canis_lupus_  659 NQTIPFVEDNVWVSDLKPRISFNFSVTSPGNVSDIIPRTEVEEAIRMYRSRINDVFRLDD
Mus_musculus  660 NQTVPFLEEDVRVSDLKPRVSFYFFVTSPQNVSDVIPRSEVEDAIRMSRGRINDVFGLND
Equus_caball  660 NQTILFGEEDVWVSDLKPRISFNFFVTSPKNASDIIPRTDVEEAIRMSRSRINDAFRLDD
Equus_asinus  638 NQTILFGEEDVWVSDLKPRISFNFFVTSPKNASDIIPRTDVEEAIRMSRSRINDAFRLDD
Capra_hircus  659 NETIPFGEENVWVSDKKPRISFKFFVTSPNNVSDIIPRTEVENAIRLCRDRINDAFQLDD
Ovis_aries__  659 NETIPFGEENVWVSDKKPRISFKFFVTSPNNVSDIIPRTEVENAIRLCRDRINDAFQLDD
Sus_scrofa__  660 NETIPFGAVDVWVSDLKPRISFNFFVTSPANMSDIIPRSDVEKAISMSRSRINDAFRLDD
Bos_taurus__  659 NETVLFGEDNVWVSDKKPRISFKFFVTSPNNVSDIIPRTEVENAIRLSRDRINDVFQLDD
Gallus_gallu  660 EQNVDFQVTDIHVGEETQRVSFYLTVSMPGNVSDIVPRADVEKAIRMSRGRISEAFRLDD
Macaca_fasci  660 HQTILFGEEDVRVADLKPRISFNFYVTAPKNVSDIIPRTEVEEAIRISRSRINDAFRLND
Macaca_mulat  660 HQTILFGEEDVRVADLKPRISFNFYVTAPKNVSDIIPRTEVEEAIRISRSRINDAFRLND
Manis_javani  660 KQTIPFEDECVRVSDLKPRVSFIFFVTLPKNVSAVIPRAEVEEAIRISRSRINDAFRLDD
Cricetulus_g  660 TQTVLFGVEDIRVSDLKPRVSFNFFVTSPQNVSDIIPRNEVEEAVRFSRGRINDVFGLDD
Mustela_puto  660 NQTIPFVGKDVRVSDLKPRISFNFIVTSPENMSDIIPRADVEEAIRKSRGRINDAFRLDD
Rhinolophus_  660 HQTILFGAENVWVSNLKPRISFNFHVTSPGNLSDIIPRPEVEGAIRMSRSRINDAFRLDD
```

```
Homo_sapiens  720 NSLEFLGIQPTLGPPNQPPVSIWLIVFGVVMGVIVVGIVILIFTGIRDRKKKNKARSGEN
Felis_catus_  720 NSLEFLGIQPTLSPPYQPPVTIWLIVFGVVMGVVVVGIVLLIVSGIRNRRKNNQARSEEN
Oryctolagus_  720 NSLEFVGIQPTLEPPYESPVPIWLVVFGVVMGMIVIGIVVLIFTGIKDRRKQKQAKREEN
Mesocricetus  720 NSLEFLGINPTLSPPYQPPVTIWLIIFGVVMGIVVVGIIILIFTGIKGRKKKNETKREEN
Canis_lupus_  719 NSLEFLGIQPTPGPPYEPPVTIWLIVFGVVMGVVVVGIVLLIFSGIRNRRKNDQARGEEN
Mus_musculus  720 NSLEFLGIHPTLEPPYQPPVTIWLIIFGVVMALVVVGIIILIVTGIKGRKKKNETKREEN
Equus_caball  720 NTLEFLGIQPTLGPPYQPPVTVWLIAFGVVMGLVVVGIVVLIATGIRGRRKKNQARSEEN
Equus_asinus  698 NTLEFLGIQPTLGPPYQPPVTVWLIAFGVVMGLVVVGIVVLIVTGIRGRRKKNQARSEEN
Capra_hircus  719 NSLEFLGIQPTLRPPYEPPVTIWLIIFGVVMGVVVIGIVVLIFTGIRDQRKKNQASSEEN
Ovis_aries__  719 NSLEFLGIQPTLRPPYEPPVTIWLIIFGVVMGVVVIGIVVLIFTGIRDQRKKNQASSEEN
Sus_scrofa__  720 NTLEFLGIQPTLGPPDEPPVTVWLIIFGVVMGLVVVGIVVLIFTGIRDRRKKKQASSEEN
Bos_taurus__  719 NSLEFLGIQPTLGPPYEPPVTIWLIIFGVVMGVVVIGIVVLIFTGIRNRRKHDNDGLQND
Gallus_gallu  720 NTLEFDGIVPTLATPYKPPVTIWLILFGVVMSLIVIGVIVLIITGQRDKRKKARGRANEA
Macaca_fasci  720 NSLEFLGIQTTLAPPYQSPVTTWLIVFGVVMGVIVAGIVVLIFTGIRDRKKKNQARSEEN
Macaca_mulat  720 NSLEFLGIQTTLAPPYQSPVTTWLIVFGVVMGVIVAGIVVLIFTGIRDRKKKNQARSEEN
Manis_javani  720 NSLEFLGIQPTLQPPYQPPVTIWLIVFGVVMGVVVVGIVVLIFTGIRDRKKKDQARSEQN
Cricetulus_g  720 NSLEFLGINPTLAPPYQPPVTIWLIIFGVVMGIVVVGIVILIVTGIRARKKNNEAKREEN
Mustela_puto  720 NSLEFLGIQPTLEPPYQPPVTIWLIVFGVVMGVVVVGIFLLIFSGIRNRRKNNQARSEEN
Rhinolophus_  720 NSLEFLGIQPTLGPPYQPPVTIWLIVFGVVMAVVVVGIVVLIITGIRDRRKTDQARSEEN

Homo_sapiens  780 PYASID-------ISKGE--NNPGFQNTDDVQTSF
Felis_catus_  780 PYASVD-------LSKGE--NNPGFQHADDVQTSF
Oryctolagus_  780 PYGFVD-------MSKGE--NNSGFQNSDDIQTSF
Mesocricetus  780 PYDSVD-------IGKGE--SNAGFLSNDDAQTSF
Canis_lupus_  779 PYASVD-------LSKGE--NNPGFQSGDDVQTSF
Mus_musculus  780 PYDSMD-------IGKGE--SNAGFQNSDDAQTSF
Equus_caball  780 PYASVD-------LSKGE--NNPGFQNGDDVQTSF
Equus_asinus  758 PYASVD-------LSKGE--NNPGFQNGDDVQTSF
Capra_hircus  779 PYGSV-------DLNKGE--NNSGFQNTDDVQTSL
Ovis_aries__  779 PYGSV-------DLNKGE--NNSGFQNTDDVQTSL
Sus_scrofa__  780 PYGSMD-------LSKGE--SNSGFQNGDDIQTSF
Bos_taurus__  779 ENLRVQQQAVKVDISRNSLKAPVPFSNSHEKLK--
Gallus_gallu  780 GSN------CEVNPYDEDGRSNKGFEQSEETQTSF
Macaca_fasci  780 PYASID-------INKGE--NNPGFQNTDDVQTSF
Macaca_mulat  780 PYASID-------INKGE--NNPGFQNTDDVQTSF
Manis_javani  780 PYASVD-------LSKGE--NNPGFQNVDDVQTSF
Cricetulus_g  780 PYDSVD-------IGKGE--SNAGFQSNDDVQTSF
Mustela_puto  780 PYASVD-------LSKGE--NNPGFQNVDDVQTSF
Rhinolophus_  780 PYSSVD-------LSKGE--NNPGFQNGDDVQTSF
```

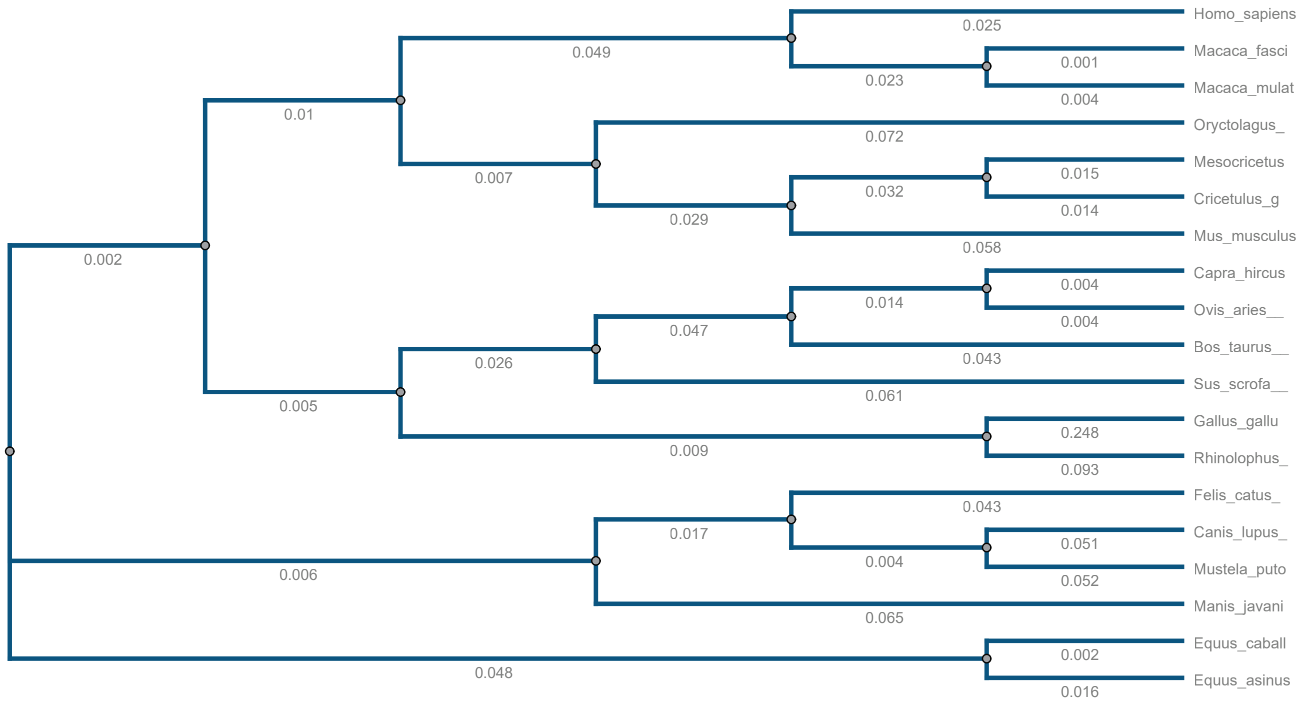


Supplemental Figure 2

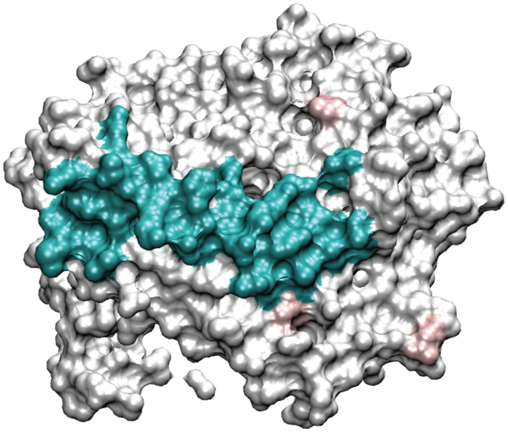

*Macaca mulatta*
*(Rhesus monkey)*

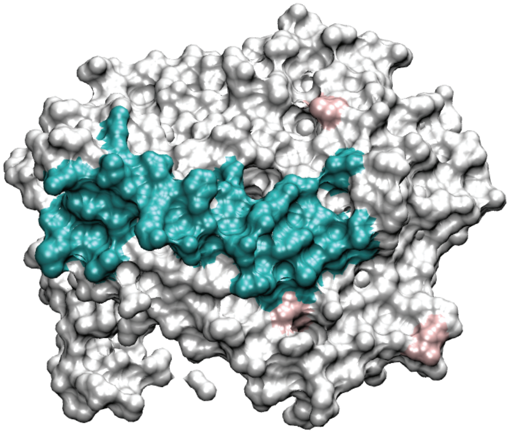

*Macaca fascicularis*
*(Cynomologous monkey)*

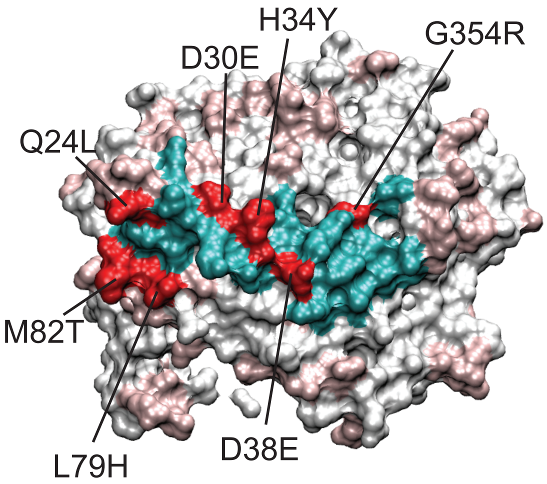


*Mustela putoris* (ferret)

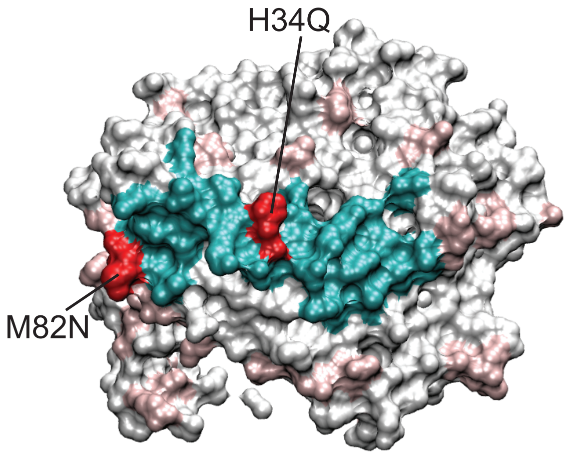


*Cricetulus griseus* (Chinese hamster)

Supplemental Figure 3

| | | *Homo sapiens* | *Macaca fascicularis* | *Macaca mulatta* | *Felis catus* | *Manis javanica* | *Cricetulus griseus* | *Canis lupus* | *Mustela putorius* | *Sus scrofa* | *Rhinolophus sinicus* | *Bos taurus* |
|---|---|---|---|---|---|---|---|---|---|---|---|---|
| human | *Homo sapiens* | 100 | 95.03 | 94.78 | 85.22 | 84.72 | 84.35 | 83.33 | 82.48 | 81.37 | 80.75 | 78.8 |
| cynomolgus monkey | *Macaca fascicularis* | 95.03 | 100 | 99.5 | 84.72 | 85.22 | 84.6 | 83.46 | 82.48 | 81.12 | 80.75 | 79.05 |
| rhesus monkey | *Macaca mulatta* | 94.78 | 99.5 | 100 | 84.47 | 84.97 | 84.35 | 83.21 | 82.36 | 80.87 | 80.5 | 78.8 |
| cat | *Felis catus* | 85.22 | 84.72 | 84.47 | 100 | 87.33 | 83.11 | 90.67 | 89.69 | 83.98 | 83.73 | 80.67 |
| pangolin | *Manis javanica* | 84.72 | 85.22 | 84.97 | 87.33 | 100 | 84.35 | 86.32 | 86.58 | 82.61 | 82.86 | 79.43 |
| chinese hamster | *Cricetulus griseus* | 84.35 | 84.6 | 84.35 | 83.11 | 84.35 | 100 | 82.71 | 82.48 | 80.5 | 80 | 79.8 |
| dog | *Canis lupus* | 83.33 | 83.46 | 83.21 | 90.67 | 86.32 | 82.71 | 100 | 89.68 | 82.09 | 81.34 | 80.02 |
| ferret | *Mustela_putorius* | 82.48 | 82.48 | 82.36 | 89.69 | 86.58 | 82.48 | 89.68 | 100 | 83.85 | 80.75 | 78.8 |
| pig | *Sus scrofa* | 81.37 | 81.12 | 80.87 | 83.98 | 82.61 | 80.5 | 82.09 | 83.85 | 100 | 80 | 84.79 |
| bat | *Rhinolophus sinicus* | 80.75 | 80.75 | 80.5 | 83.73 | 82.86 | 80 | 81.34 | 80.75 | 80 | 100 | 77.56 |
| cow | *Bos taurus* | 78.8 | 79.05 | 78.8 | 80.67 | 79.43 | 79.8 | 80.02 | 78.8 | 84.79 | 77.56 | 100 |

**Supplemental Figure 4**

**Supplemental Figure 4.**

**A. Binding curves for immobilized ACE2-Fc binding RBD analyte.**

The RBD binding responses to both ACE2 proteins shows a surface density dependence. As the ACE2 surface density increases, the off rate of bound RBD slows down. The figures below show an overlay of the normalized response for each complex. The higher the density surface the lower the noise sturcture on the data and the slow the apparent dissociation rate. For example, for the Human ACE2 (1 uM RBD) data set the data from the lowest to highest density surface goes red, green, blue and black (3 fold titration up to 1.0 $\mu$M). If the binding interaction was simple the different colored responses would overlay with one another. However, a trend is seen in the data indicating that a more stable complex is formed at higher density surfaces. This is often an indication that the analyte (in this case RBD) is multimerizing producing an avidity effect on the surface.

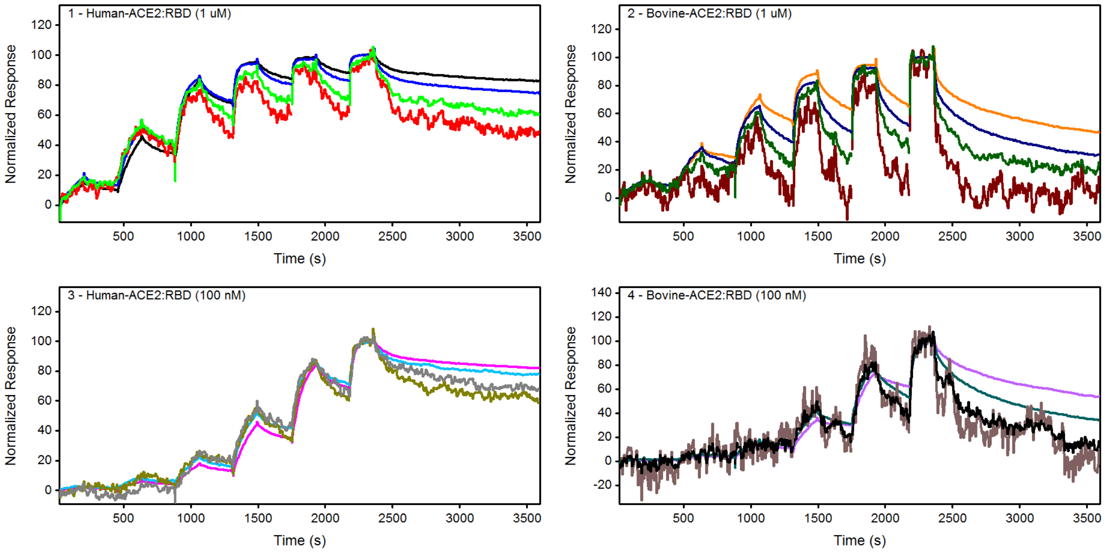

### B. Two-site model

The data for each data set were fit to a two independent site model in order to assign values to the interactions. A summary of the binding constants is provided in the table below.

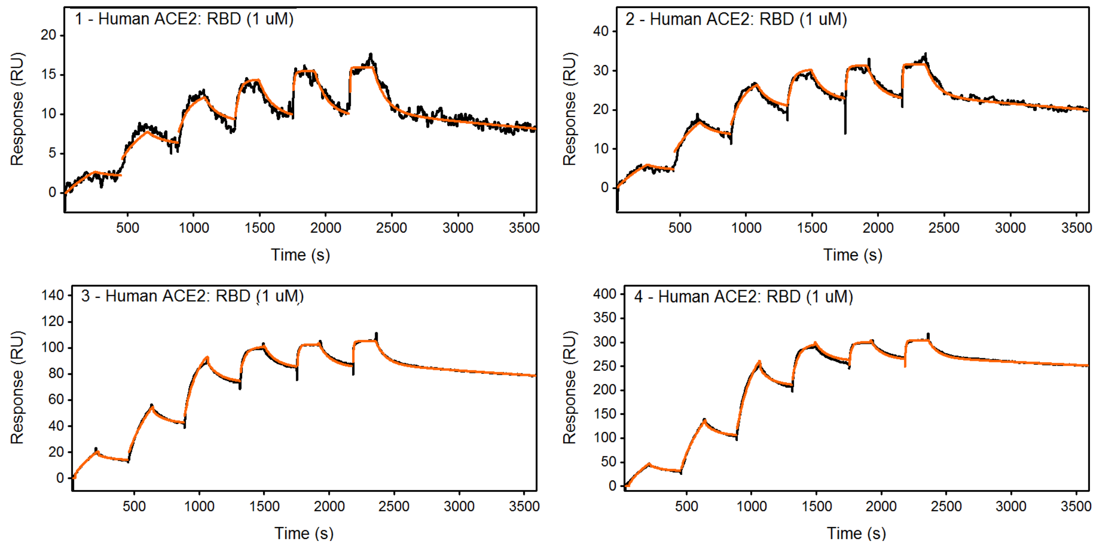


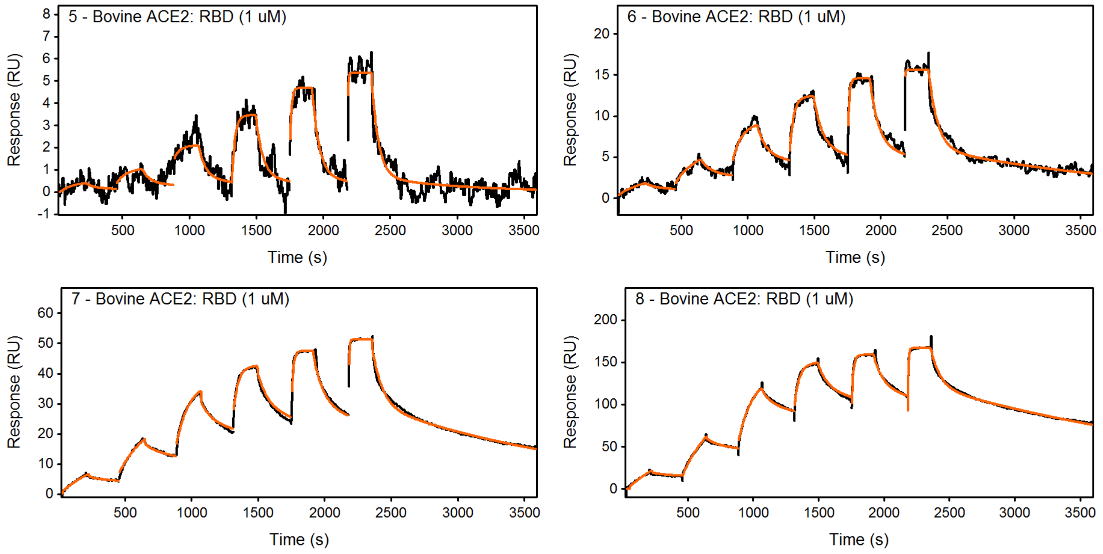

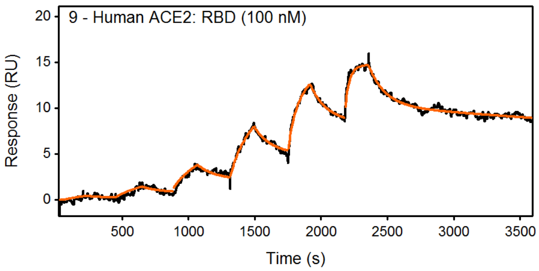
9 - Human ACE2: RBD (100 nM)
Response (RU)
Time (s)

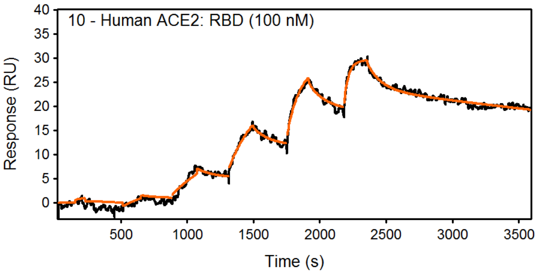
10 - Human ACE2: RBD (100 nM)
Response (RU)
Time (s)

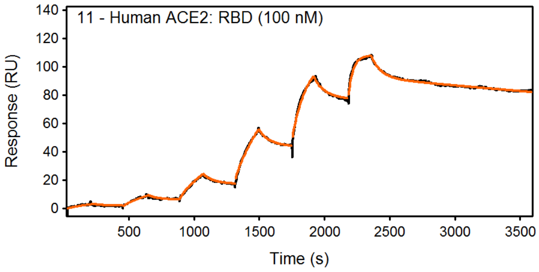
11 - Human ACE2: RBD (100 nM)
Response (RU)
Time (s)

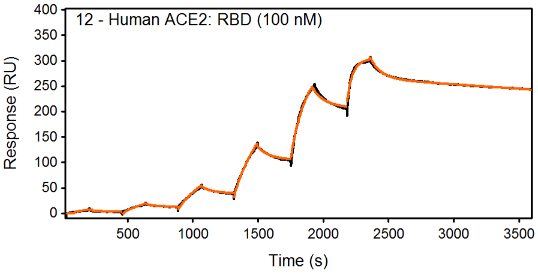
12 - Human ACE2: RBD (100 nM)
Response (RU)
Time (s)

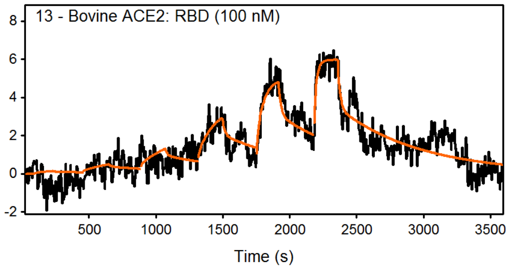
13 - Bovine ACE2: RBD (100 nM)
Time (s)

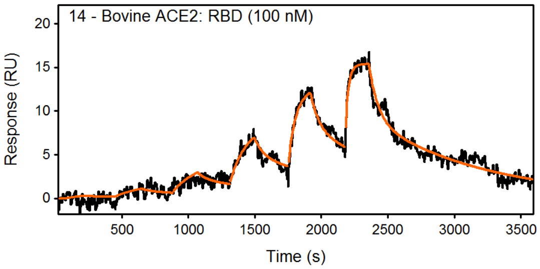
14 - Bovine ACE2: RBD (100 nM)
Response (RU)
Time (s)

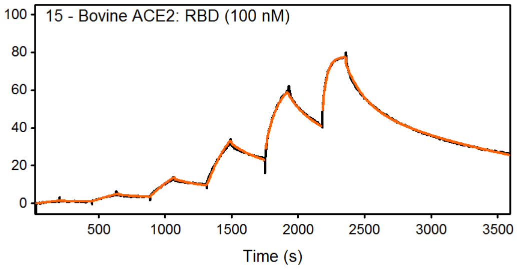
15 - Bovine ACE2: RBD (100 nM)
Time (s)

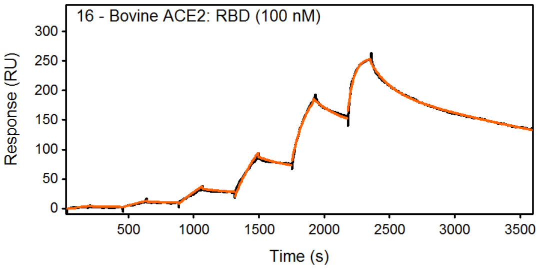
16 - Bovine ACE2: RBD (100 nM)
Response (RU)
Time (s)

#### C. Binding constants determined for two site model.

| ACE2 surface | Highest [RBD] | $R_{max1}$ (RU) | $k_{a1}$ ($M^{-1}s^{-1}$) | $k_{d1}$ ($s^{-1}$) | $K_{D1}$ (nM) | $R_{max2}$ (RU) | $k_{a2}$ ($M^{-1}s^{-1}$) | $k_{d2}$ ($s^{-1}$) | $K_{D2}$ (nM) |
|---|---|---|---|---|---|---|---|---|---|
| Human | 1 uM | 6.9 | 1.50E5 | 0.011 | 75 | 10 | 3.80E5 | 1.90E-4 | 0.5 |
| Human | 1 uM | 10 | 2.00E5 | 0.012 | 59 | 24 | 3.11E5 | 1.30E-4 | 0.4 |
| Human | 1 uM | 18 | 8.00E5 | 0.012 | 15 | 87 | 2.41E5 | 8.40E-5 | 0.3 |
| Human | 1 uM | 39 | 1.30E6 | 0.013 | 10 | 268 | 1.95E5 | 5.20E-5 | 0.3 |
| Human | 100 nM | 4.8 | 1.13E6 | 0.008 | 7 | 10 | 2.94E5 | 1.10E-4 | 0.4 |
| Human | 100 nM | 5.9 | 1.10E6 | 0.009 | 8 | 24 | 2.83E5 | 1.60E-4 | 0.6 |
| Human | 100 nM | 16 | 2.50E6 | 0.013 | 5 | 93 | 2.70E5 | 9.40E-5 | 0.3 |
| Human | 100 nM | 47 | 1.20E6 | 0.012 | 10 | 267 | 2.30E5 | 6.90E-5 | 0.3 |

| | | | | | | | | | |
|---|---|---|---|---|---|---|---|---|---|
| Bovine | 1 uM | 5.1 | 2.10E5 | 0.019 | 90 | 0.7 | 5.00E5 | 1.60E-3 | 3.2 |
| Bovine | 1 uM | 10 | 2.30E5 | 0.015 | 66 | 6.1 | 2.70E5 | 5.90E-4 | 2.2 |
| Bovine | 1 uM | 20 | 1.60E5 | 0.013 | 82 | 30 | 2.36E5 | 5.56E-4 | 2.4 |
| Bovine | 1 uM | 48 | 1.90E5 | 0.014 | 72 | 121 | 2.20E5 | 3.78E-4 | 1.7 |
| Bovine | 100 nM | 3.2 | 2.00E6 | 0.060 | 30 | 3.6 | 4.90E5 | 1.63E-3 | 3.3 |
| Bovine | 100 nM | 8.1 | 4.80E5 | 0.015 | 31 | 10 | 3.70E5 | 1.17E-3 | 3.2 |
| Bovine | 100 nM | 29 | 2.30E5 | 0.007 | 29 | 53 | 2.90E5 | 5.70E-4 | 2.0 |
| Bovine | 100 nM | 77 | 1.20E5 | 0.007 | 55 | 203 | 2.22E5 | 3.27E-4 | 1.5 |
